# Genetic modifiers of somatic expansion and clinical phenotypes in Huntington’s disease reveal shared and tissue-specific effects

**DOI:** 10.1101/2024.06.10.597797

**Authors:** Genetic Modifiers of Huntington’s Disease (GeM-HD) Consortium, Jong-Min Lee, Zachariah L. McLean, Kevin Correia, Jun Wan Shin, Sujin Lee, Jae-Hyun Jang, Yukyeong Lee, Kyung-Hee Kim, Doo Eun Choi, Jeffrey D. Long, Diane Lucente, Ihn Sik Seong, Ricardo Mouro Pinto, James V. Giordano, Jayalakshmi S. Mysore, Jacqueline Siciliano, Emanuela Elezi, Jayla Ruliera, Tammy Gillis, Vanessa C. Wheeler, Marcy E. MacDonald, James F. Gusella, Anna Gatseva, Marc Ciosi, Vilija Lomeikaite, Hossameldin Loay, Darren G. Monckton, Christopher Wills, Thomas H. Massey, Lesley Jones, Peter Holmans, Seung Kwak, Cristina Sampaio, Michael Orth, G. Bernhard Landwehrmeyer, the European Huntington’s Disease Network (EHDN) Registry investigators, Jane S. Paulsen, the Huntington Study Group (HSG) PREDICT-HD investigators, E. Ray Dorsey, the HSG COHORT investigators PHAROS and TREND-HD investigators, Richard H. Myers, the HD-MAPS investigators

## Abstract

Huntington’s disease (HD), due to expansion of a CAG repeat in *HTT*, is representative of a growing number of disorders involving somatically unstable short tandem repeats. We find that overlapping and distinct genetic modifiers of clinical landmarks and somatic expansion in blood DNA reveal an underlying complexity and cell-type specificity to the mismatch repair-related processes that influence disease timing. Differential capture of non-DNA-repair gene modifiers by multiple measures of cognitive and motor dysfunction argues additionally for cell-type specificity of pathogenic processes. Beyond *trans* modifiers, differential effects are also illustrated at *HTT* by a 5’-UTR variant that promotes somatic expansion in blood without influencing clinical HD, while, even after correcting for uninterrupted CAG length, a synonymous sequence change at the end of the CAG repeat dramatically hastens onset of motor signs without increasing somatic expansion. Our findings are directly relevant to therapeutic suppression of somatic expansion in HD and related disorders and provide a route to define the individual neuronal cell types that contribute to different HD clinical phenotypes.

## INTRODUCTION

Genes in which mutations underlie Mendelian disorders have typically been found through molecular genetic strategies that uncover genetic variants of strong effect, thereby starting the search for molecular mechanisms and targets for intervention. One of the earliest such genes identified, *HTT*, underlying Huntington’s disease (HD [MIM: 143100]), was found three decades ago (Huntington’s Disease Collaborative Research Group, 1993), but this knowledge has yet to lead to an effective therapy. HD involves progressive neurodegeneration resulting in characteristic motor abnormalities, cognitive decline, and psychiatric manifestations caused by an expanded (> 35) CAG trinucleotide repeat in *HTT* that lengthens a polyglutamine tract in the protein huntingtin (Huntington’s Disease Collaborative Research Group, 1993). To date, clinical trial attempts to treat HD by reducing huntingtin expression have not succeeded.

We have demonstrated that the genome-wide association study (GWAS) strategy using genetic variants common in the general population, typically used to identify risk loci in complex diseases, can also be applied to discover modifiers of Mendelian disease. Characterization of CAG repeat lengths and adjacent sequences in the HD population (Ciosi et al., 2019; Cubo et al., 2019; Lee et al., 2012; Wright et al., 2019) and GWAS exclusively of HD participants (Genetic Modifiers of Huntington’s Disease Consortium, 2015, 2019; Lee et al., 2022) have shown that: 1) HD individuals with two expanded *HTT* alleles display an age at onset of motor signs (i.e., motor onset) consistent with complete dominance of the longer of the two CAG repeats; 2) the length of the uninterrupted inherited CAG repeat is the primary determinant of the timing with which the characteristic HD motor signs emerge; and, 3) the timing of motor onset is not due to the length of the inherited huntingtin polyglutamine tract, which can differ despite identical uninterrupted CAG repeats due to glutamine-encoding CAA codons that distinguish canonical and non-canonical repeat regions. Tellingly, the ages at motor onset and other clinical landmarks are influenced by genetic variation at DNA-maintenance genes, implicating length-dependent somatic expansion of the *HTT* CAG repeat in driving the rate of pathogenesis. Somatic expansion of the *HTT* CAG repeat is observed in post-mortem HD brain (Kennedy et al., 2003; Matlik et al., 2024; Pressl et al., 2024), with the longest repeats in those individuals who showed the earliest onset of disease symptoms (Swami et al., 2009). Together, these genetic findings support a model of HD pathogenesis comprising sequential components: an initial phase in which the inherited HD-associated CAG repeat undergoes further expansion in vulnerable neurons, leading, when it has reached a sufficient length, to a consequent phase where the cells are unable to cope with the resultant toxicity (Hong et al., 2021). This domino scenario produces progressive dysfunction, neuropathology, and clinical manifestations, culminating in the early death of the individual.

Recently, single-cell analysis of post-mortem HD brains has supported this model and proposed that the CAG expansion phase can be subdivided at a CAG length (∼80 CAGs) beyond which the rate of expansion accelerates dramatically (Handsaker et al., 2024). HD genetic analysis, therefore, points to distinct therapeutic options: 1) preventing or slowing CAG expansion; 2) raising the threshold at which CAG expansion accelerates; 3) altering the CAG repeat length threshold that triggers toxicity; 4) interfering with the toxicity mechanism; or 5) alleviating the consequences of ongoing neuronal loss. However, neither the mechanism by which the expanded repeat is further expanded nor how the somatically-expanded CAG repeat drives cellular toxicity is known with any certainty. GWAS-identified modifiers that are DNA-maintenance genes imply some type of DNA ‘mismatch repair’ and those that are not DNA-maintenance genes do not appear to converge on a single mechanism. Cellular damage could potentially be triggered at the DNA, RNA or protein levels, including polyglutamine-induced toxicity.

To further inform understanding of the mechanisms involved in HD pathogenesis and provide direct insights into the molecular mechanism of somatic expansion, we increased our clinical sample size to > 16,000 HD individuals, expanded the GWAS to assess multiple clinical landmark phenotypes and performed a distinct GWAS of *HTT* CAG somatic expansion measured directly in blood DNA. New participants were sequenced to determine uninterrupted CAG repeat length and power assessment of adjacent sequences. We find that CAG repeat expansion is driven by a mismatch repair-related process whose modification and consequences show unexpected complexity, including different participants and effects in different cell types. In addition to identifying new modifier loci acting on CAG expansion, we report new loci that act differentially on clinical phenotypes and are not obviously related to DNA repair. Notably, an *HTT* 5’-UTR sequence variant upstream from some canonical CAG repeats is associated with enhanced CAG expansion in blood DNA but does not influence clinical landmarks. Remarkably, non-canonical alleles with loss of a synonymous CAACAGCCGCCA sequence at the end of the repeat strongly hasten motor onset without enhancing CAG expansion, suggesting that these alleles accelerate HD at another step in pathogenesis. Our findings are directly relevant to the therapeutic suppression of somatic repeat expansion in HD and related disorders and lay the foundation for dissecting HD pathogenesis by relating phenotypic outcomes to cell-type specific genetic modification to uncover the cell populations underlying different features of the disorder.

## RESULTS

### GWAS of HD motor onset and algorithmically predicted age at clinical landmarks

Focusing on HD age at motor onset (defined as the age at which motor signs emerged as determined by an expert rater) as a single point in the lifelong trajectory of HD, we previously reported a GWAS of 9,058 individuals of European ancestry whose genome-wide SNV (small nucleotide variant – including both single and multi-nucleotide variants) data were imputed from array-typed genotypes using data from the Haplotype Reference Consortium (HRC) (Genetic Modifiers of Huntington’s Disease Consortium, 2019). These analyses revealed that inherited uninterrupted CAG repeat length is the strongest single predictor of age at motor onset. Subsequently, we re-imputed these genotypes using the Trans-Omics for Precision Medicine (TOPMed) Program data as a reference panel for GWAS and considered two additional clinically defined disease landmarks from the Unified Huntington’s Disease Rating Scale (UHDRS): 1) a Diagnostic Confidence Level of 4 (DCL4) indicating > 99% confidence that motor abnormalities are unequivocal signs of HD; and, 2) a score of 6 on the 13 point Total Functional Capacity scale (TFC6), corresponding to a substantial reduction in the capacity to function (a score of 13 reflects normal, full functioning in the judgement of a trained rater) (Lee et al., 2022). TFC6 reflects a marginal engagement in one’s occupation, needing major assistance to handle financial affairs, impaired capacity to manage domestic responsibilities, and at least mild impairment of ability to perform activities of daily living (Shoulson and Fahn, 1979). The ages at which a participant reached these clinical landmarks, reflecting the diagnostic appearance and significant deterioration of HD manifestations, respectively, were predicted algorithmically by integrating available UHDRS clinical measures (Lee et al., 2022). Age at DCL4 and TFC6 proved to be informative, highly correlated phenotypes (Pearson correlation coefficient 0.97) in a GWAS of < 7,000 HD participants.

To begin to examine genetic modification of HD across the entire disease trajectory, we have now increased our overall sample size to 16,399 HD individuals, including 12,982 with a recorded age at motor onset and 11,698 with sufficient clinical data to have been included in the previous algorithmic prediction of age at TFC6. Most of the additional individuals were deeply phenotyped participants from the Enroll-HD platform (Langbehn et al., 2023; Sathe et al., 2021). To maximize the sample size in each type of analysis, we included 513 individuals genotyped in this and prior studies who did not meet the criteria for European ancestry. We initially examined the same phase of the HD trajectory as in the previous GWAS with this expanded participant sample. Figure 1A displays the GWAS results at minor allele frequency (MAF) > 1%, illustrating the increased power obtained with algorithmically predicted age at TFC6 over rater-determined age at motor onset. As expected, age at DCL4 (n = 11,408) produced very similar results (Figure S1A). Neither limiting the GWAS to participants of European ancestry (Figure S1B) nor, disregarding infrequent SNVs (MAF < 1%), with the important exception of *HTT* (discussed below), substantially altered the pattern of the results (Figure S1C & D). The expanded GWAS analyses identified several new significant signals at two new loci and at a previously reported locus (magenta genes in Figure 1A), adding to the repertoire of HD modifiers while reproducing at greater significance the previously reported pattern for six loci encompassing DNA-maintenance genes (*PMS1, MLH1, MSH3, PMS2, FAN1, LIG1*) and three loci harboring other candidate genes (*TCERG1, RRM2B, CCDC82*). New loci included an additional gene involved in DNA maintenance, *POLD1,* and a regulator of gene expression, *MED15*.

**Figure 1.**
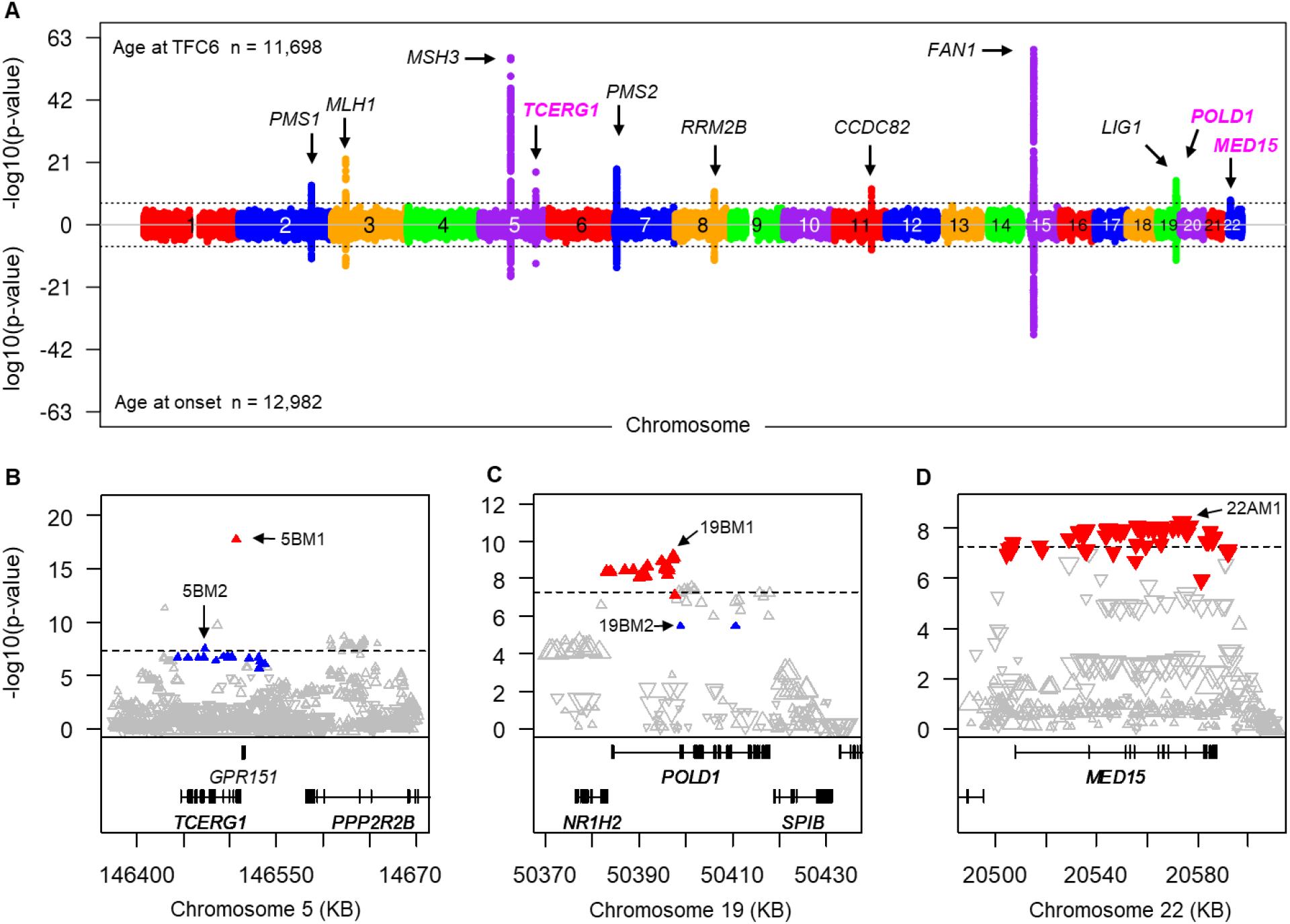
– The age at TFC6 clinical landmark detects new modifier effects. **A.** GWAS results are plotted with each point representing a test SNV at > 1% MAF and significance shown as -log10(p-value) (top) for age at TFC6 and log10(p-value) (bottom) for age at motor onset. The dashed black line represents the threshold for genome-wide significance (p = 5.0E-08). Loci with new signals are shown in colored font. Inflation factors were 1.010 for age at motor onset and 0.997 for age at TFC6. See also Figures S1 and S2 and Table S1. **B**-**D.** Association results for age at TFC6 from panel A are shown in the *TCERG1* region (**B**), the *POLD1* region (**C**) and the *MED15* region (**D**), with genes in each region shown below the plot. For each distinguishable modifier effect, the top SNV and all other SNVs showing r^2^ >0.8 with the top SNV are shown as filled triangles (either red for 5BM1, 19BM1 and 22AM1, or blue for 5BM2 and 19BM2) while all other SNVs are represented by unfilled triangles. The size of each triangle reflects the SNV MAF, which can be judged by comparison with the triangles representing top tag SNVs for distinguishable modifier effects in Table S1 (e.g., MAF: 5BM1 2.7%; 5BM2 2.0%; 19BM1 9.7%; 19BM2 1.0%; 22AM1 21.3%). The triangle orientation reflects the direction of effect (upward pointing = delaying; downward pointing = hastening).

We also performed pathway analysis of the GWAS results using gene-wide p-values for all 18,919 genes. When tested for pathway enrichment, age at motor onset, age at TFC6 and age at DCL4 association signals detected gene sets involving DNA mismatch repair as the only Bonferroni significant results (based on correction for 10,043 pathways tested), consistent with the timing of these phenotypes being driven by the somatic expansion of the *HTT* CAG repeat. When the significant DNA-maintenance genes (*FAN1*, *MSH3, PMS1, PMS2, MLH1, LIG1* and *POLD1*) were excluded from the analysis to focus on pathways that could reflect aspects of the subsequent cellular toxicity mechanism, the top results, GO:0032371 “regulation of sterol transport” and GO:0032374 “regulation of cholesterol transport” (both p = 7.7E-06 with age at TFC6) and GO:0010644 “cell communication by electrical coupling (p = 2.1E-05 with age at motor onset), did not survive Bonferroni correction.

### Many modifier loci display multiple distinguishable effects

The functional impact of variation at a modifier locus can reflect the presence of multiple contributing variants, including sequences not captured directly as SNVs, that may act in concert or separately. Indeed, we previously reported the existence of more than one distinguishable modifier effect at each of *PMS1, PMS2, MSH3, FAN1, and LIG1* and adopted a nomenclature consisting of the chromosome number, the order of locus discovery on that chromosome (i.e., A, B, etc.), and a sequential number for each distinguishable modifier effect at that locus (i.e., M1, M2, M3, etc.). Consequently, in our expanded data set, we defined and named distinguishable modifier effects at newly discovered loci and revisited the modifier effects at previously reported loci using a series of sequential conditional analyses based on effect size and significance (Table S1).

At *TCERG1*, which encodes a regulator of transcriptional elongation and pre-mRNA splicing (Figure 1B), conditional analysis disclosed a new second modifier effect (5BM2). In this and subsequent analyses, filled colored triangles depict the top tag SNV for a distinguishable modifier effect along with those SNVs showing r^2^ > 0.8 with that top SNV. The up- or down-pointing orientation of the triangle indicates the direction of effect, and the triangle size reflects the SNV MAF. We previously described a delaying age at motor onset modifier effect (5BM1), tagged by rs79727797, and subsequently attributed modification to the length of a quasi-tandem hexamer repeat encoding a glutamine/alanine tract in TCERG1 (Lobanov et al., 2022). This SNV tag (red in Figure 1B) showed the greatest significance (age at motor onset: p=8.9E-14; age at TFC6: p=1.8E-18), but conditional analyses in the age at TFC6 GWAS revealed a second modifier haplotype of significant SNVs spanning *TCERG1* (blue triangles in Figure 1B). This distinct modifier effect (designated 5BM2; top SNV rs202157262) was genome-wide significant for delayed age at TFC6 (p = 3.3E-08) but did not achieve suggestive significance for age at motor onset (p = 9.3E-05). Conditioning the age at TFC6 analysis on either of the two 5BM1 or 5BM2 tag SNVs increased the significance for the other, and simultaneously conditioning on both SNVs eliminated all suggestive significant signals. The 5BM2 modifier is not associated with length variation in the *TCERG1* hexamer repeat, being present only on chromosomes with the most frequent repeat length (91% of alleles) in the exome data used previously to analyze 5BM1 (Lobanov et al., 2022). 5BM2 tag SNVs do not reveal significant expression quantitative trait locus (eQTL) or splicing quantitative trait locus (sQTL) signals in brain in the GTEx database (https://www.gtexportal.org/home/aboutAdultGtex).

Among the newly discovered loci, a genome-wide significant landmark-delaying modifier effect (19BM1; top SNV rs112837068; red triangles in Figure 1C) was detected at *POLD1* by age at TFC6 (p = 6.6E-10), but not by age at motor onset (p = 5.1E-04). A second landmark-delaying effect (19BM2; blue triangles in Figure 1C) was tagged at suggestive significance for both age at TFC6 and age at motor onset and (p = 3.4E-06 and 3.1E-06, respectively) by the *POLD1* missense variant, rs3218772 (Arg30Trp). Conditioning the age at TFC6 analysis on either the top 19BM1 or 19BM2 tag SNV increased the significance for the other and conditioning on them together removed all suggestive significant SNV signals. The 19BM1 and 19BM2 tag SNVs do not show significant brain eQTLs or sQTLs in GTEx for any coding gene in the region, although the former is associated with the expression of *NAPSB,* a transcribed pseudogene (e.g., p = 8.6E-08 in putamen and 3.9E-07 in frontal cortex in GTEx). POLD1 has both DNA polymerase and 3’ to 5’ exonuclease activity and is involved in DNA replication and repair (Nicolas et al., 2016), making it likely that this locus influences the somatic expansion phase of HD pathogenesis.

A new landmark-hastening modifier (22AM1) at *MED15* (top SNVs rs177425 and rs165666; red triangles in Figure 1D) on chromosome 22 yielded genome-wide significance with age at TFC6 (p = 5.0E-09) but not even nominal significance with age at motor onset (p = 7.7E-02), again reinforcing the differential informativeness at some loci for the algorithmically predicted disease landmark compared with clinical rater-determined age at motor onset.

Conditioning the age at TFC6 analysis on rs177425 removed all suggestive significant signals, supporting a single modifier effect. SNVs tagging the 22AM1 modifier effect do not display significant brain eQTLs or sQTLs in GTEx. *MED15* specifies a subunit of the Mediator complex involved in the regulated transcription of RNA polymerase II-dependent genes and is required for cholesterol-dependent gene regulation (Nakatsubo et al., 2014; Yang et al., 2006). This subunit localizes to the Mediator tail that bridges enhancers and promotors, making it a candidate for modification of HD pathogenesis through a mechanism other than promoting *HTT* CAG expansion. Interestingly, *MED15* contains a glutamine-rich region encoded in part by a polymorphic CAG repeat whose predominant allele comprises 12 CAGs (Sandhu et al., 2004). The rs177425 minor allele is in linkage disequilibrium with the 13 CAG allele (r^2^ = 0.40, D’=0.63), suggesting that this repeat could be involved in the modifier effect.

At *PMS1, PMS2, MSH3, FAN1, and LIG1,* the conditional analyses confirmed multiple distinguishable modifier effects (Figure S2) while *MLH1*, *RRM2B*, and *CCDC82* loci continued to exhibit single modifier effects (Table S1).

### Non-canonical synonymous *HTT* CAG repeat sequences are age at motor onset modifiers

To overcome the potential inaccuracies in determining the *HTT* CAG repeat length by fragment-sizing, for this GWAS we determined the uninterrupted CAG repeat length by paired-end MiSeq sequencing across the repeat and adjacent sequence for all new participants and those available from prior studies (n = 7,553). Most HD chromosomes carried the canonical sequence (CAG)_n_CAACAGCCGCCA(CCG)_n_ but we also confirmed four primary alternative non-canonical sequences on disease chromosomes (Ciosi et al., 2019; Genetic Modifiers of Huntington’s Disease Consortium, 2019; McAllister et al., 2022): CAACAG duplication, (CAG)_n_(CAACAG)_2_CCGCCA(CCG)_n_, here called CAACAG-dup; CAA loss also missing the CCA triplet, (CAG)_n_(CCG)_n_, here called CAA/CCA-loss; CAA loss only, (CAG)_n_CCGCCA(CCG)_n_, here called CAA-loss; and, CCA-loss only, (CAG)_n_CAACAG(CCG)_n_, here called CCA-loss (Table 1).

**Table 1.**
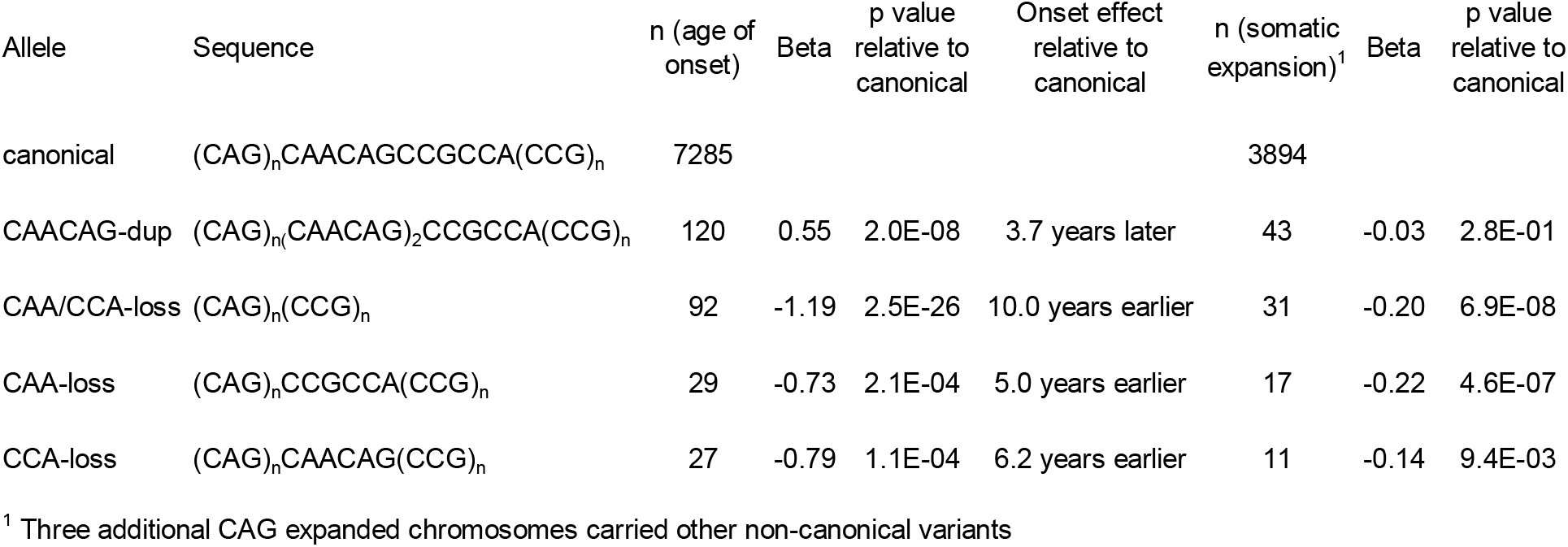
Association of non-canonical HTT CAG repeat-adjacent sequences with age at motor onset and somatic expansion

SNV-based association analysis with age at motor onset, carried out at MAF > 0.1% and correcting for uninterrupted repeat length, revealed significant signals only for infrequent SNVs (MAF < 1%) in the terminal 5 Mb of chromosome 4p containing *HTT* and conditional analyses determined that these SNVs tag haplotypes bearing two of the non-canonical alleles, CAA/CCA-loss and CAACAG-dup (Figure S3), but not CAA-loss or CCA-loss. However, examination of all CAA/CCA-loss carriers revealed that some did not carry the tag SNVs making them imperfect proxies for this non-canonical sequence.

We also tested the individual non-canonical sequences on the expanded CAG chromosome directly for association with age at motor onset (Table 1), again correcting for the uninterrupted CAG length (the use of fragment length-based CAG determinations in the natural history studies used for algorithmic predictions prevented accurate testing of other disease landmarks). The CAA/CCA-loss allele (n = 92) was strongly associated with earlier motor onset (p = 2.5E-26) than canonical alleles with the same uninterrupted CAG repeat lengths.

Interestingly, those participants with the CAA/CCA-loss sequence who lacked the SNV-tagged haplotype (n = 31 of the 92) showed a similar effect on age at motor onset to those with the shared haplotype (10.0 versus 10.4 years earlier than expected), arguing that the modifier influence is due directly to the CAA/CCA-loss sequence rather than a linked modifier. Although there were fewer carriers of expanded alleles with only CAA-loss or CCA-loss (n = 29 and 27, respectively), these also showed associations (p = 2.1E-04 and 1.1E-04) with hastening of motor onset (5.0 and 6.2 years). In contrast, the CAACAG-dup allele (n = 120) on CAG expanded chromosomes was genome-wide significant (p = 2.0E-08) for later than expected motor onset (3.7 years). Notably, canonical alleles show variation downstream of the CAG repeat region in the form of a polymorphic CCG repeat. Most canonical HD chromosomes (n = 7,285) carry 7 CCGs (n = 6,827) with 10 CCGs (n = 400) being the next most frequent.

However, unlike the non-canonical variations, this polymorphism on canonical HD alleles showed no association with age at motor onset (p = 0.24). Thus, the association data implicate a direct influence of the CAA/CCA-loss, and probably the other non-canonical sequences, on modifying age at motor onset. However, they do not distinguish whether these variants alter the instability of the adjacent CAG repeat in the first phase of HD pathogenesis or have some other effect.

### Motor onset-hastening CAA/CCA-loss alleles do not increase *HTT* CAG repeat expansion

To address directly whether the non-canonical *HTT* repeat sequences influence CAG repeat instability, we turned to blood as an accessible source of somatic DNA that shows sufficient repeat expansion for genetic analysis (Ciosi et al., 2019). We focused on a subset of 3,999 GWAS participants where MiSeq sequencing had been performed using a protocol optimized to capture variation in repeat expansion along with the repeat-adjacent sequences (Ciosi et al., 2018). For each participant, most of whom carried canonical alleles (n = 3,894), we calculated a ‘somatic expansion ratio’ (SER) as the number of uninterrupted CAG reads longer by one to ten CAGs than modal length divided by the number of uninterrupted CAG reads of modal length. The SER revealed an age- and length-dependent increase in CAG expansion against which the non-canonical alleles could be compared (Figure 2A). Counterintuitively, the CAA/CCA-loss allele, tied strongly to the hastening of HD motor onset with the expectation of increased somatic expansion, was associated in blood DNA with a significant reduction in the SER (p versus canonical = 6.9.E-08) (Figure 2B). The CAA-loss allele was also associated with reduced expansion (p versus canonical = 4.6E-07), while the CCA-loss and the CAACAG-dup alleles showed less evidence of an effect (p versus canonical = 9.4E-03 and 2.8E-01, respectively, both for reduced expansion).

**Figure 2.**
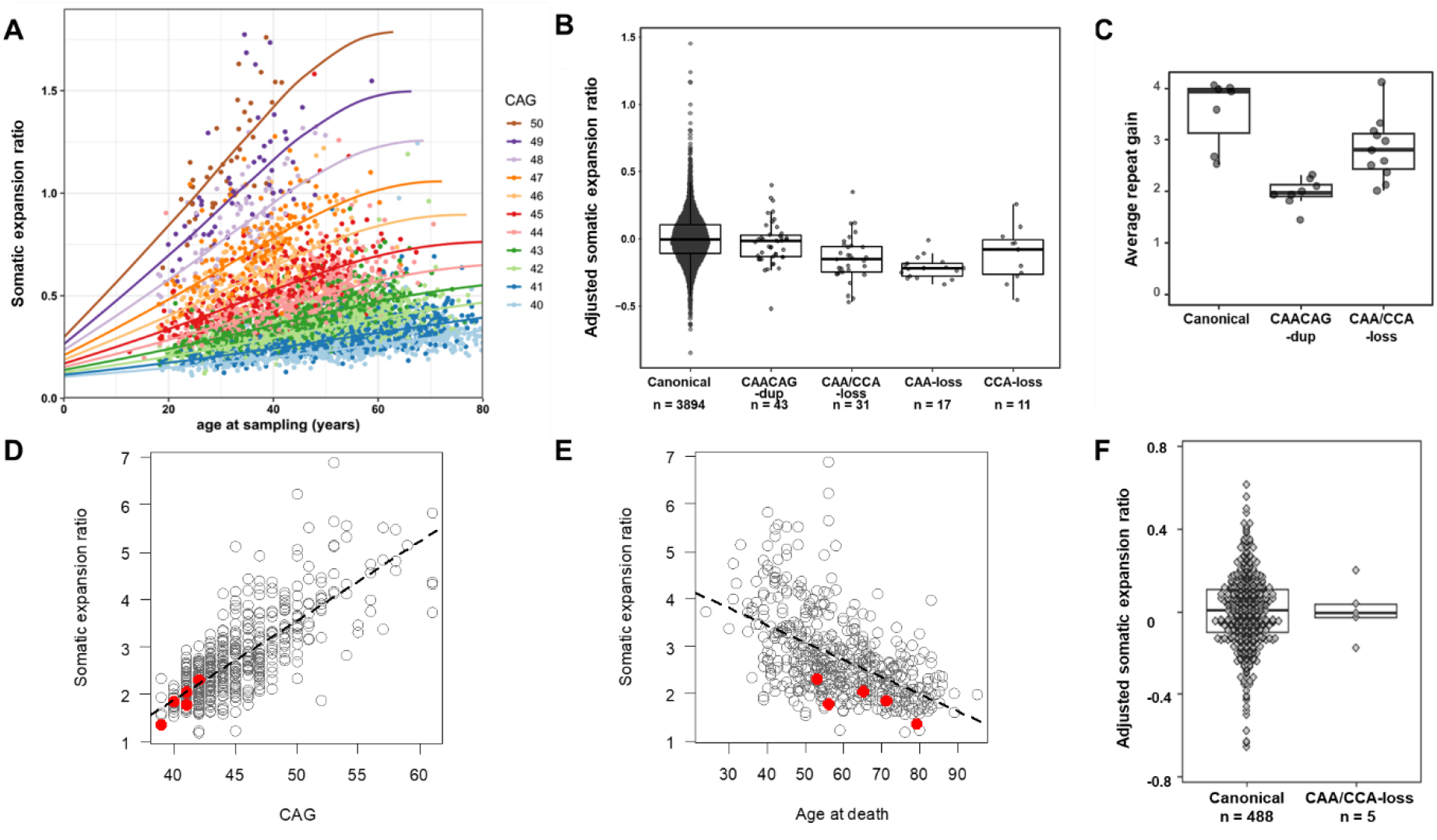
– Non-canonical alleles do not increase the rate of CAG expansion. **A.** The somatic expansion ratio of the *HTT* exon 1 CAG repeat in blood DNA is allele length- and age-dependent. The scatterplot shows the somatic expansion ratio (SER = 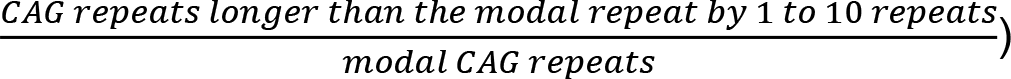 in blood DNA plotted against age at sampling in 3,999 Enroll-HD participants. Points are color-coded based on the length of the inherited uninterrupted CAG length determined by MiSeq. Lines on the scatterplot are the SER values predicted by the multiple linear regression ln(SER) ∼ *β*_0_ + *β*_1_.CAG + *β*_2_.age + *β*_3_.(CAG x age) + *β*_4_.CAG^2^ + *β*_5_.age^2^ + *β*_6_.(CAG^2^ x age) + *β*_7_.(CAG x age^2^) + *ε*_0_. Boxplot (25th and 75th percentiles (box), median (line), and range (whiskers, capped at 1.5x the interquartile range)) of the relationship between canonical and non-canonical *HTT* repeat sequences with the somatic expansion ratio in blood DNA adjusted for CAG length, age, CCG length and PCR batch (*i.e.,* the residuals of the multiple linear regression ln(SER) ∼ β_0_ + β_1_.CAG + β_2_.age + β_3_.(CAG x age) + β_4_.CAG^2^ + β_5_.age^2^ + β_6_.(CAG^2^ x age) + β_7_.(CAG x age^2^) + β_8_.CCG + β_9_.PCRbatch). **B.** Boxplot of average number of CAG repeats gained over four weeks for *HTT* exon 1 variants knocked into the AAVS1 locus in RPE1 cells. Clonal knock-in RPE1 lines with canonical (n = 7), CAACAG-dup (n = 8), or CAA/CCA-loss (n = 11) *HTT* exon 1 variants are each represented by a single dot, positioned based on the average repeat gain of triplicate cultures. The average repeat gain represents the difference between mean CAG repeat (weighted on the fragment analysis peak height) at four-weeks and day-zero, which was adjusted for the effect size of day zero mean repeat length that ranged between 112-120 CAGs. Both CAACAG-dup and CAA/CCA-loss alleles showed significantly reduced CAG repeat expansion compared to canonical (p < 0.001 and = 0.015, respectively). **C.** Somatic expansion ratio length in post-mortem frontal cortex from 488 HD individuals carrying canonical expanded repeat sequences (open circles) and 5 carrying CAA/CCA-loss HD alleles (filled red circles) is plotted versus modal *HTT* CAG repeat **D.** Somatic expansion ratio in post-mortem frontal cortex from 488 HD individuals carrying canonical repeat sequences (open circles) and 5 carrying CAA/CCA-loss HD chromosomes (filled red circles) is plotted versus their ages at death. **E.** Boxplot of the relationship between canonical and non-canonical *HTT* repeat sequences with the somatic expansion ratio in HD frontal cortex DNA, adjusted for CAG length, age, CCG length and PCR batch as described in B. The CAA/CCA-loss allele carriers were not significantly different from those with canonical alleles (p = 0.96)

To further examine the relationship of the two most frequent non-canonical sequences and CAG repeat expansion, we capitalized on human hTERT-immortalized retinal pigment epithelial cells (RPE1), found recently to support CAG repeat expansion when maintained in confluent cultures (McLean et al., 2024). We created three types of lines carrying canonical, CAA/CCA-loss, or CAACAG-dup *HTT* exon 1 with 112-120 uninterrupted CAG repeats knocked into the AAVS1 safe-harbor locus. After four weeks at confluency, a series of clones with the canonical sequence showed a median gain of 3.9 CAG repeats while the CAACAG-dup and CAA/CCA-loss clones gained fewer repeats (Figure 2C), at 2.0 and 2.8 CAGs (p versus canonical < 0.001 and = 0.015, respectively, accounting for starting average CAG repeat length). Although reduced CAG expansion in the RPE1 cells due to the adjacent CAACAG-dup sequence is consistent with its association with HD clinical delay, we saw no significant effect for the CAACAG-dup allele in the blood DNA expansion ratio. However, neither the RPE1 cells nor the blood DNA instability assay suggested that the CAA/CCA-loss sequence promotes greater expansion of the adjacent CAG repeat.

The decade earlier-than-expected motor onset of HD individuals with a CAA/CCA-loss allele cannot be explained by an inherent increase in propensity for expansion, presenting two alternatives: either this non-canonical allele behaves differently with respect to CAG repeat expansion in blood versus brain, or its clinical modifier impact is not through directly increasing somatic expansion. Consequently, we used paired-end MiSeq sequencing to screen HD post-mortem frontal cortex samples for canonical and non-canonical alleles. Only the CAA/CCA-loss allele yielded multiple samples (five) for comparison with 488 carrying a canonical sequence. The canonical sample SERs revealed a CAG length-dependent increase from which the CAA/CCA-loss samples did not appear to deviate (Figure 2D). When compared by age at death, the CAA/CCA-loss allele carriers had an SER at the low end of the range of canonical allele carriers with the same lifespan (Figure 2E). Finally, when both uninterrupted CAG repeat length and age at death were controlled for, the adjusted SERs for CAA/CCA-loss and canonical alleles were not different (Figure 2F). Thus, like the blood and cell line studies, these post-mortem brain data provided no evidence for an increase in somatic expansion due to the non-canonical CAA/CCA-sequence, though such a difference would have been expected from the highly significant association of these alleles with age at motor onset.

### GWAS identifies modifiers of CAG repeat expansion in blood DNA

To identify loci that influence somatic expansion, we carried out a GWAS for this molecular phenotype measured in blood DNA. Seeking maximum sample size, we initially explored a set of 5,342 DNA sizing traces collected from standard PCR fragment-length genotyping of study participants. In an approach analogous to the SER, we assessed CAG expansion using the Peak Proportional Sum (PPS) method, which takes the ratio of summed peak heights longer than the modal CAG length divided by the height of the modal repeat signal. Correcting for CAG length, age at collection, and their interaction term, we performed a PPS GWAS that showed a robust signal at *HTT* on chromosome 4, along with genome-wide significant signals on chromosomes 2, 5, 14, 15, and 17 for SNVs with MAF > 1% (Figure S4A). However, matching to the subset of participants with available sequence data revealed that the *HTT* region signal was due to a confounding artefact of the standard genotyping assay: due to mis-priming the CAACAG-dup sequence consistently produces a distinctive PCR fragment ‘shoulder’ signature that scores as highly significant repeat expansion but does not reflect biological instability (Figure S4B). Consequently, we used the 3,999 participants with MiSeq SER data to perform a GWAS at MAF > 1%, correcting for CAG length, age, CCG length, PCR batch and CAG repeat-adjacent sequence. This GWAS yielded more robust peaks at the same loci as the PPS GWAS, along with an additional genome-wide significant peak on chromosome 7 (Figure 4A, Table S2). For the somatic expansion modifier effects, we adopted a nomenclature similar to that for clinical modifier effects consisting of chromosome number with a letter noting the order of locus discovery on that chromosome (all ‘A’ at present), followed by the designator ‘BE’ for blood expansion, and a sequential number for each distinguishable modifier effect at that locus (i.e., M1, M2, M3, etc.).

Interestingly, we also saw a strong genome-wide significant peak on chromosome 4 where the minor allele of the most significant SNV, rs146151652 (p = 4.8E-30), was associated with increased somatic expansion. This SNV (upward pink triangle in Figure 4B) is located in the 5’-UTR of *HTT*, upstream from a canonical repeat sequence. Conditioning the MiSeq blood DNA expansion GWAS on rs146151652 left no SNVs at suggestive significance in the vicinity of *HTT*, indicating a single modifier effect (4ABEM1). Notably, rs146151652 also emerged atop the peak remaining when we conditioned the PPS association analysis mentioned above to remove the effects of the CAACAG-dup allele artefact (p = 8.2E-16; Figure S4C). In a computationally phased analysis of these PPS data, comparing individuals imputed to be heterozygous and homozygous for the major allele at rs146151652, the modifier effect was driven by those with the rs146151652 minor allele in *cis* with the expanded repeat (n = 205, p = 1.2E-14) versus in *trans*, n = 118, p = 0.0068). Notably, rs146151652 is not associated with the clinical landmarks (p > 5.0E-02), suggesting that it is not tied to increased somatic expansion in HD target neurons in the brain. This 5’-UTR SNV yields a significant *cis*-eQTL for increased *HTT* expression (p = 3.9E-16) in CD16+ neutrophils in the BLUEPRINT project (Chen et al., 2016), but only for *HTT-AS1* (p = 5.1E-11) and *GRK4* (p = 4.1E-08) in the more extensive eQTLGen data (Vosa et al., 2021).

At the mismatch repair protein gene *MSH2* on chromosome 2, a modifier haplotype, 2ABEM1, was associated with decreased CAG expansion (downward pink triangles in Figure 3C). The top SNV (rs113983130, p = 3.7E-42) is not associated with *MSH2* expression in blood, while many highly significant *MSH2* eQTLs did not show evidence for modification of CAG expansion. However, whilst we have quantified somatic expansion in circulating white blood cells, the age-dependent nature of somatic expansion suggests that expansions accrue in hematopoietic stem cells. Unfortunately, hematopoietic stem cells are not represented in available eQTL datasets and the relationship between eQTL observed in circulating blood cells and hematopoietic stem cells primarily resident in the bone marrow remains unknown. Interestingly, the 2ABEM1 effect is also tagged by a missense variant, rs4987188 (Gly322Asp; p = 5.3E-40), associated with the intergenerational transmission of *de novo* changes in the lengths of dinucleotide and tetranucleotide but not trinucleotide repeats (Kristmundsdottir et al., 2023). Conditioning on rs113983130 revealed a second modifier effect, 2ABEM2 (teal triangles in Figure 3C), nearer to another mismatch repair protein gene, *MSH6,* ∼300 kb centromeric to *MSH2*. The minor allele of the top 2ABEM2 SNV (rs79939438, p= 1.3E-09) is associated with increased *HTT* CAG repeat expansion and increased expression of *MSH6* in blood (eQTLGen, p = 1.2E-24). Neither of the tags for the 2ABEM1 or 2ABEM2 modifier effects, nor any other SNVs in this region, achieved suggestive significance for an influence on age at motor onset or age at TFC6.

**Figure 3.**
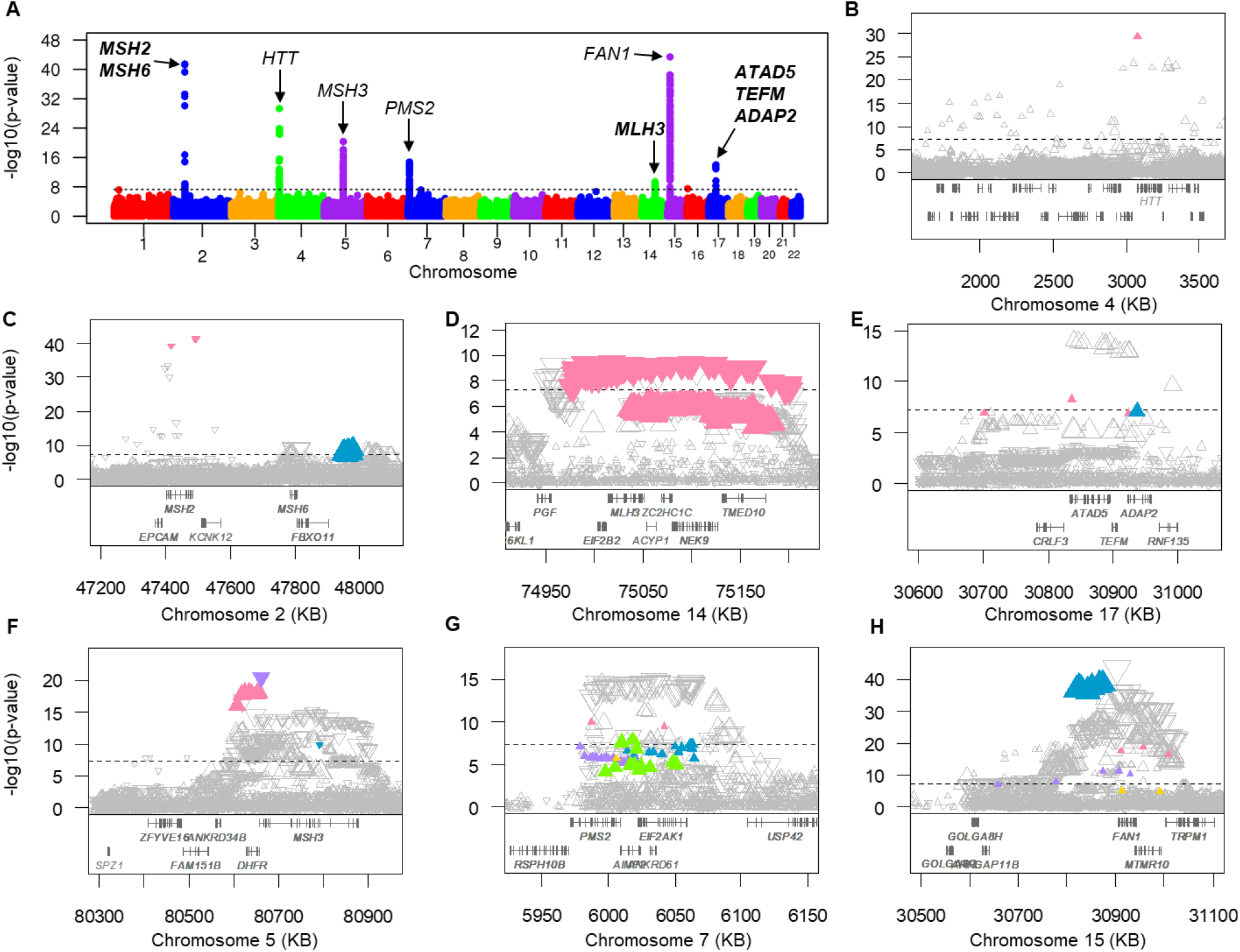
– GWAS of CAG repeat expansion measured in blood DNA. **A.** GWAS results for adjusted somatic expansion ratio determined in blood DNA are plotted with each point representing a test SNV at > 1% MAF and candidate genes labeled at each peak. The dashed black line represents the threshold for genome-wide significance (p = 5.0E-08). See also Figure S3 and Table S2. **B**-**H**. Association results for adjusted somatic expansion ratio from panel A are shown in the regions of *HTT* (**B**), *MSH2 / MSH6* (**C**), *MLH3* (**D**), *ATAD5* (**E**), *MSH3* (**F**), *PMS2* (**G**), and *FAN1* (**H**) with genes in each region shown below the plot. Each SNV is represented by a triangle whose size and orientation reflect its MAF and direction of effect (upward pointing = increased expansion; downward pointing = decreased expansion). **B**. The top SNV (filled pink triangle) tagging modifier effect 4ABEM1 is in the 5’-UTR of *HTT* and does not show r^2^ > 0.8 with any other SNV. Conditioning on the top SNV removed all significant signals. **C**. Two significant modifier effects defined by conditional analysis: pink = 2ABEM1; teal = 2ABEM2. **D**. Conditional analysis supported a single significant modifier effect: pink = 14ABEM1. **E**. Conditional analysis revealed two distinguishable modifier effects (pink = 17ABEM1; teal = 17ABEM2) that are both captured by, and account for, the most significant SNVs on the plot. **F**. Three significant modifier effects were defined by conditional analysis: pink = 5ABEM1; teal = 5ABEM2; light purple = 5ABEM3. Although the 5ABEM2 tag is an infrequent SNV with MAF < 1%, it was included here since it was previously identified as the tag for 5AM2 in the much larger clinical GWAS. **G**. Conditional analysis revealed five distinguishable modifier effects (pink = 7ABEM1; teal = 7ABEM2; light purple = 7ABEM3; gold = 7ABEM4; green = 7ABEM5) combinations of which are captured by and account for the most significant SNVs on the plot. **H**. Conditional analysis identified four distinguishable modifier effects: pink = 15ABEM1; teal = 15ABEM2; light purple = 15ABEM3; gold = 15ABEM4. The 15ABEM1 and 15ABEM3 tags are missense variants both present at < 1% MAF in this dataset but which were previously identified tags for 15AM1 and 15AM3 in the much larger clinical GWAS.

On chromosome 14 at another mismatch repair gene, *MLH3*, a single significant modifier effect (14ABEM1, pink triangles in Figure 3D) was tagged by immediately adjacent SNVs rs56375606 and rs56329719 (p = 4.1E-10; together also known as rs386778872), associated with decreased CAG expansion. Although the top SNVs are not listed as significant in eQTLGen, they are part of a large set of frequent SNVs in very high linkage disequilibrium (LD), some of which are annotated as robust eQTLs associated with reduced *MLH3* expression (e.g., rs175016, p < 3.3E-310 in blood). Unlike the *MSH2-MSH6* region, in the clinical GWAS the *MLH3* region yielded suggestive significant signals. For both age at motor onset and age at TFC6, these signals were associated with disease hastening albeit with different top SNVs (rs12879725, p = 6.3E-06 and rs175438, p = 3.7E-06, respectively).

The modifier signal on chromosome 17 is in the vicinity of three genes (Figure 3E): *ATAD5* (ATPase family AAA domain containing 5), whose product participates in the cellular DNA damage response, *TEFM* (transcription elongation factor, mitochondrial), and *ADAP2* (ArfGAP with dual PH domains 2). Conditional analyses showed that the top SNV (rs62070643) captured two distinguishable modifier effects associated with increased somatic expansion: 17ABEM1 (pink triangles in Figure 3E) tagged by rs117662433 (p = 4.91E-09) and 17ABEM2 (teal triangle in Figure 3E) tagged by rs8067252 (p = 8.50E-08). Conditioning simultaneously on both rs117662433 and rs8067252 reduced the significance of all other SNVs to below suggestive significance. None of the SNVs in this region achieved suggestive significance in the clinical GWAS.

The remaining significant loci (Figure 3F-H) from the somatic expansion GWAS implicated DNA repair genes *MSH3, PMS2* and *FAN1*, which all showed multiple modifier effects in the clinical GWAS (Table S2). Likewise, conditional analyses also revealed multiple effects at each of these loci in the somatic expansion GWAS (Figure S2). For *PMS2* and *FAN1,* where the peak SNVs in the somatic expansion GWAS showed a very frequent minor allele that appeared to be associated with decreased CAG expansion, the conditional analyses revealed that this peak was a by-product of multiple distinguishable CAG expansion-promoting effects on chromosomes with the major allele at the peak SNV. Thus, simultaneous conditioning on the tag SNVs for the multiple less frequent CAG expansion-promoting modifier effects at each locus removed all suggestive significant SNVs, including the initial top SNVs.

### Shared clinical and CAG expansion modifier loci show both coincident and distinct effects

The presence of modifier signals at *MSH3, MLH3, PMS2* and *FAN1* in both the age at TFC6 and somatic expansion GWAS indicated that variation at these loci acts on both cell types responsible for these phenotypes. The TFC6 phenotype is driven by dysfunction or loss of brain neurons contributing to motor control and cognitive function. The blood CAG expansion phenotype is presumed to reflect CAG repeat instability in hematopoietic stem cells. In Figure 4, we compare the significance and direction of effect of SNVs at *MSH3, MLH3*, *PMS2*, and *FAN1* in the age at TFC6 (left panels) and blood CAG expansion (right panels) GWAS. Variants associated with clinical hastening of HD, which is expected to involve increased CAG expansion in target neurons, are shown as downward triangles in the left panels. Variants associated with increased CAG expansion in blood DNA are shown as upward triangles in the right panels.

**Figure 4.**
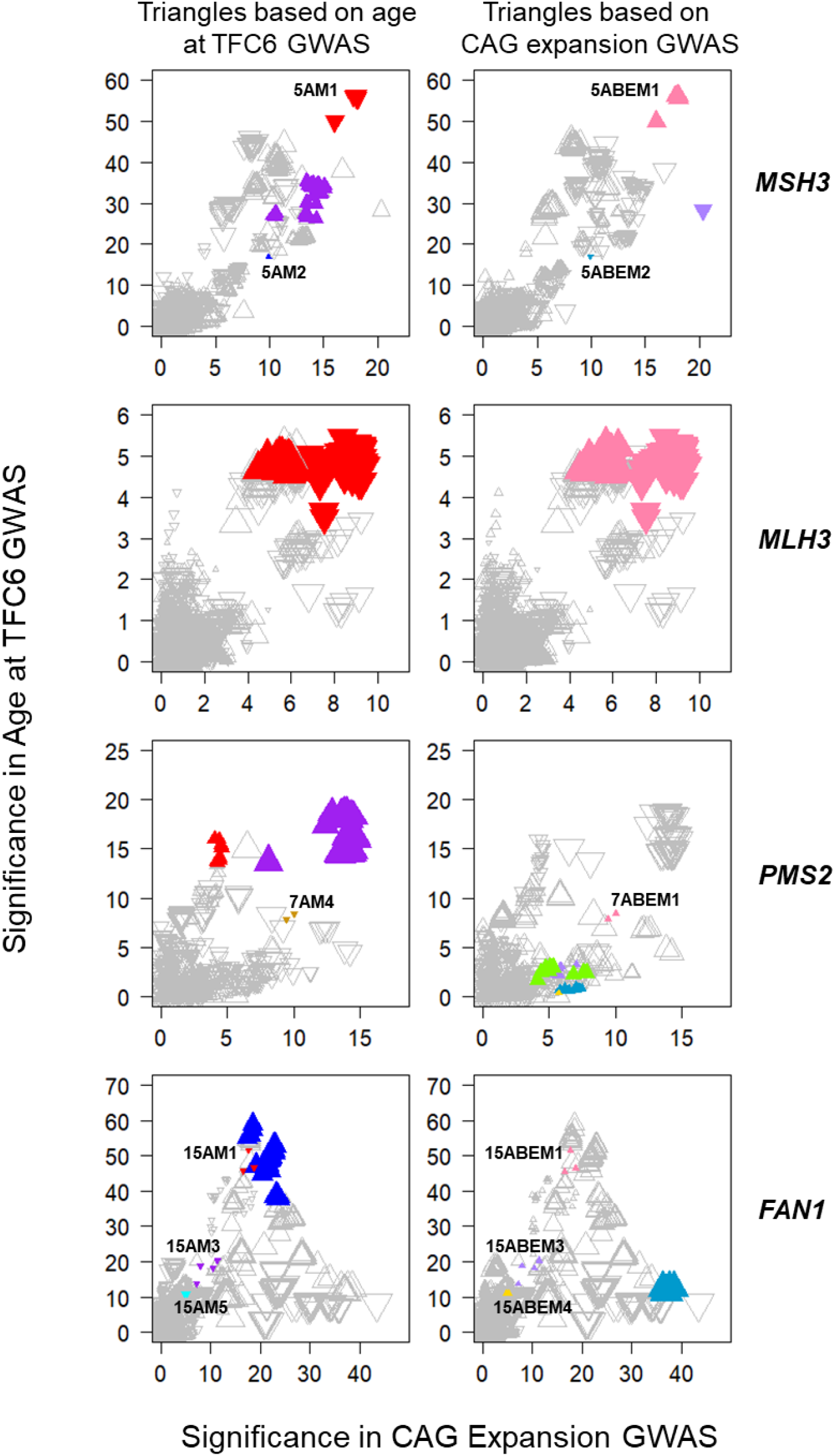
– Blood CAG expansion and age at TFC6 capture both coincident and distinct modifier effects. The significance of SNVs (plotted as -log10(p-value) in the *MSH3*, *MLH3*, *PMS2*, and *FAN1* regions is directly compared between the age at TFC6 GWAS and the blood somatic expansion GWAS. The left-hand panel of each pair plots SNVs as triangles with direction based on the age at TFC6 GWAS: upward pointing = delaying effect; downward pointing = hastening effect. The right-hand panel of each pair plots the same set of SNVs as triangles with direction based on the somatic expansion GWAS: upward pointing = increased expansion; downward pointing = decreased expansion. SNVs tagging clinical modifier effects (left panels) determined by conditional analysis are filled red, blue, purple, brown, and cyan for effects 1 through 5 respectively (see Table S1). SNVs tagging CAG expansion modifier effects (right panels) determined by conditional analysis are filled pink, teal, light purple, gold, and green for effects 1 through 5 respectively (see Table S2). Based on the model of somatic CAG expansion in target neurons driving the timing of clinical landmarks, tag SNVs associated with earlier or later motor onset would be expected to show increased or decreased CAG expansion in blood, respectively, if the same modifier mechanism acted in both tissues. Those modifier effects that appear to show such concordance are labeled in the figure. Filled triangles pointing in the same direction between paired panels or appearing in only one of the paired panels suggest instances where the molecular basis for phenotype modification differs between the brain and the blood.

The *MSH3* region has shown three significant clinical modifier effects (top left panel): a rare landmark-delaying effect (5AM2, blue triangle) and common hastening (5AM1, red triangles) and delaying (5AM3, purple triangles) effects (Table S1). The somatic expansion GWAS similarly showed two common effects (top right panel), one increasing (5ABEM1, pink triangles) and the other decreasing (5ABEM3, purple triangle) CAG expansion, along with a significant expansion-decreasing influence (5ABEM2, teal triangle) marked by the same infrequent SNV as 5AM2 (rs113361582, Tables S2 & S3). Like the 5AM2 / 5ABEM2 effects, the 5AM1 and 5ABEM1 effects also coincided, tagged by top SNVs rs245100 and rs245105 which are in almost complete LD (r^2^ = 0.998). However, the 5AM3 and 5ABEM3 modifier effects appeared distinct. The strong correspondence and agreement in the expected direction of effect for 5AM1 / 5ABEM1 and 5AM2 / 5ABEM2 tag SNVs argue that within each of these pairs, a molecular mechanism underlying modification is likely the same for the clinical landmark and blood CAG expansion. However, the distinct patterns for SNVs tagging the 5AM3 and 5ABEM3 effects suggest that these effects reflect different molecular drivers of modification by *MSH3* in the different cell types contributing to the phenotypes.

At *MLH3*, the modifier haplotype (14ABEM1, pink triangles, second right panel) is very frequent, with SNV allele frequencies close to 0.5. This haplotype carries minor alleles at some sites and major alleles at others, generating different directions of effect (relative to minor alleles), with the most robust signals showing reduced expansion in Figures 4D and 5 (second right panel). However, a similar pattern of common SNVs was associated with hastening of TFC6 (red triangles, Figure 4, second left panel), which is expected to involve increased somatic expansion in brain. Interestingly, the top 14ABEM1 SNV tags do not appear as brain eQTLs in GTEx, but instead are significant brain sQTLs (caudate, p = 1.1e-14; putamen, p = 1.5E-14; frontal cortex, p = 2.6e-10), indicating that they capture different *MLH3* regulation in blood and brain.

**Figure 5.**
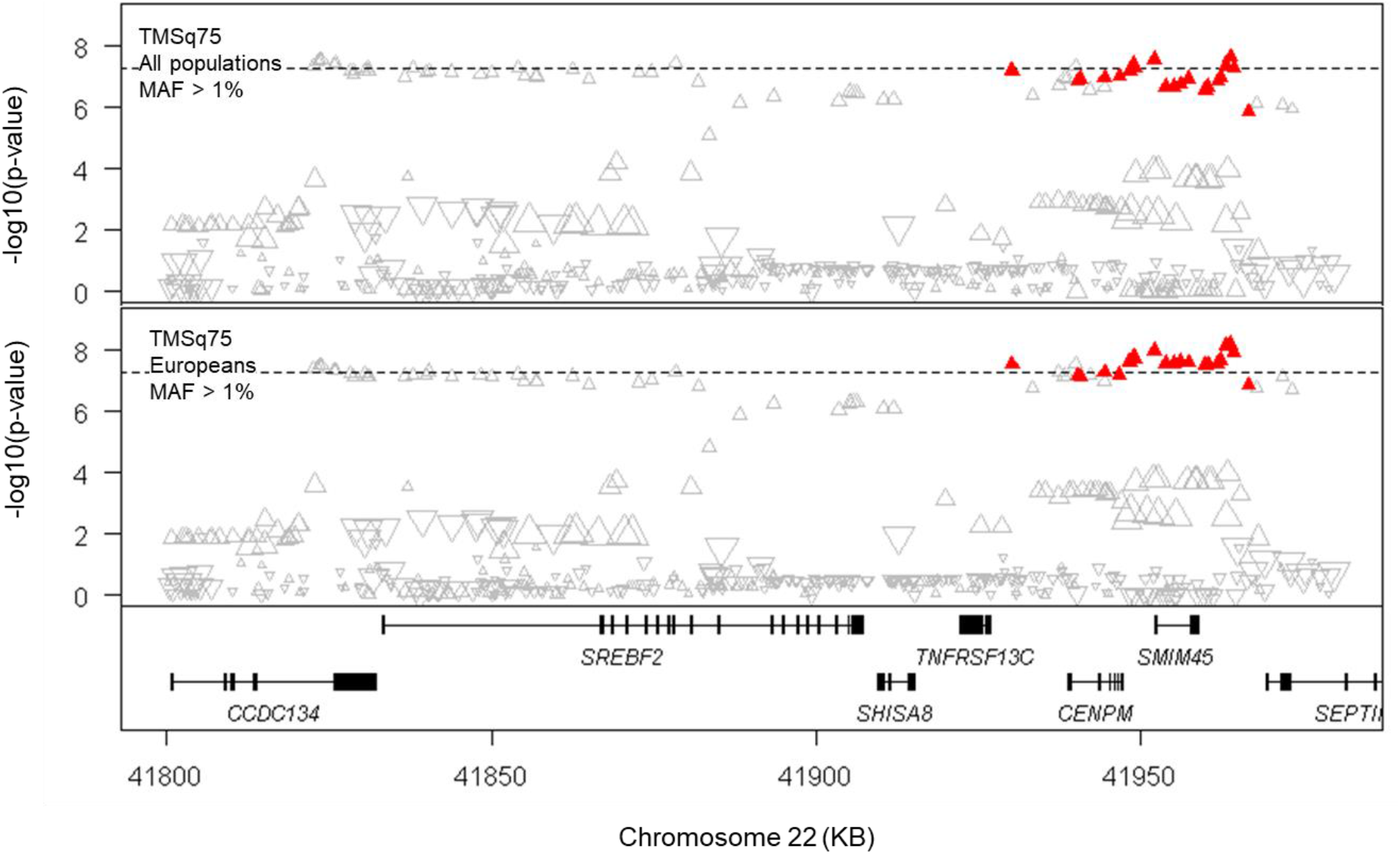
– A new modifier locus in a gene-dense region of chromosome 22. Association signals are shown at a new clinical modifier locus on chromosome 22 showing genome-wide significant signal with the algorithmically predicted late-disease motor landmark TMSq75 (see Figure S5 and Table S). The top panel and bottom panels show results from all individuals and only individuals of European ancestry, respectively, above the genes on the region. The dashed black line represents the threshold for genome-wide significance (p = 5.0E-08). The top SNV and all other SNVs showing r^2^ > 0.8 with the top SNV are shown as filled red triangles while all others are shown as unfilled triangles. The size and orientation of each triangle reflects its MAF and direction of effect (upward pointing = delaying; downward pointing = hastening). Conditional analysis based on the top SNV removed all significant signals.

At *PMS2*, sequential conditional analyses in our expanded dataset revealed three distinguishable age at TFC6 modifier effects (Table S1), with the previously reported 7AM2 effect no longer showing suggestive significance when analysis was conditioned simultaneously on 7AM1, 7AM3 and 7AM4. Conditional analysis of the somatic expansion GWAS data showed five distinguishable effects associated with increased CAG expansion. Only one of these, 7ABEM1 (Figure 4, pink triangles, third right panel), is tagged by the same SNV (rs1805324, a Met662Ile variant reported to affect PMS2-MLH1 interaction (Yuan et al., 2002)) as a clinical modifier effect, 7AM4 (Figure 4, brown triangles, third left panel). The 7ABEM1 effect (increased somatic instability) and 7AM4 effect (hastening of TFC6) are consistent with the same mechanism operating on the blood and brain phenotypes. Notably, the 7ABEM4 tag is also a missense variant (rs63750123, Ile18Val) but shows no signal in the clinical GWAS (p = 0.41). The lack of correspondence of 7AM1 and 7AM3 with 7ABEM2, 7ABEM3, 7ABEM4, and 7ABEM5 suggests that these distinguishable effects reflect differentially modifiable regulation of *PMS2* in the brain and hematopoietic stem cells.

The *FAN1* region also showed partial correspondence between the modifier effects in the clinical and somatic expansion GWAS. Three clinical hastening effects, 15AM1, 15AM3 and 15AM5 (Figure 4, bottom left panel, red, purple and cyan triangles), the first two of which are tagged by missense variants that affect FAN1 function (Kim et al., 2020; McAllister et al., 2022), were directly matched (bottom right panel, pink, light purple and gold triangles) by 15ABEM1 (rs150393409, Arg507His), 15ABEM3 (rs151322829, Arg377Trp) and 15ABEM4 (rs118089305), associated with increased CAG expansion. However, the common 15ABEM2 effect (teal triangles, bottom right panel) did not correspond to the common landmark-delaying 15AM2 (blue triangles, bottom left panel) but revealed a distinct pattern of SNVs associated with increased somatic expansion. Interestingly, the 15ABEM2 tag SNP, rs61260387, is an eQTL whose minor allele is associated with increased expression of *FAN1* in blood. This seeming contradiction with FAN1’s CAG expansion-suppressing activity likely indicates that 15ABEM2 exerts differential effects on the circulating cells tested in expression analysis and the hematopoietic stem cells in which age-dependent CAG expansions must accumulate.

### Looking later in HD identifies a new HD clinical modifier

The GWAS hits from somatic expansion in blood DNA and those from clinical landmarks were only partially overlapping, indicating similarities and differences in the influences on somatic expansion in different tissues. Although some of the modifier effects are associated with coding sequence changes, the majority are likely to involve regulatory differences. This suggests that modification of CAG expansion might also differ due to regulatory difference between different brain cell types, and the same could be true of the consequent neuronal toxicity mechanism(s). Indeed, we have previously reported that age landmarks early in the DCL4-TFC6 interval based on motor (UHDRS Total Motor Score (TMS)) or cognitive (Symbol Digit Modalities Test (SDMT)) measures capture both similarities and differences in modifier signals (Lee et al., 2022). To increase the opportunity to detect and compare somatic instability and toxicity modifiers by lengthening the time interval over which they could act, we initially used the TMS and SDMT measures, which respectively increase and decrease with HD progression, to generate age landmark phenotypes later in the disease process. We created late landmark phenotypes by algorithmically predicting each individual’s age at the 75% quantile (TMSq75) or 25% quantile (SDMTq25) value of all data points in the observational studies that contributed to our dataset. The ages at these landmarks differ by inherited CAG length, but as a comparison, for individuals inheriting 42 uninterrupted CAGs, the average ages at TMSq75 and SDMTq25 were 63.4 and 65.1, versus 48.0 and 57.8 at DCL4 and TFC6.

GWAS of the age at TMSq75 (n = 11,683) but not the age at SDMTq25 (n = n = 11,324) phenotype revealed a new genome-wide significant modifier locus (top SNP rs8137409, p = 2.3E-08) on chromosome 22 (Figure 5 top, see also Figure S5 and Table S1). This new landmark-delaying effect, 22BM1, maps to a gene-dense region with several candidates, including *CCDC134* (coiled-coil domain containing 134), *SREBF2* (sterol regulatory element binding transcription factor 2), *SHISA8* (shisa family member 8), *TNFRSF13C* (TNF receptor superfamily member 13C), *CENPM* (centromere protein M), *SMIM45* (small integral membrane protein 45), and *SEPTIN3* (a member of the septin family of GTPases). Conditional analysis pointed to a single modifier effect tagged most efficiently by rs8137409 and rs59906307, which flank *SREBF2* (whose product physically interacts with MED15) at either end of a broad haplotype. However, these two SNVs only displayed moderate linkage disequilibrium (r^2^ = 0.57 in our dataset). Consequently, we examined the region using only the participants of European origin, and the peak signal became more significant (p = 6.2E-09) at rs8137409 (Figure 5 bottom). The lack of a known DNA repair gene in this vicinity suggests that the 22BM1 modifier might act through a gene in this region to influence HD pathogenesis after initial somatic repeat expansion.

### Late trajectory HD clinical phenotypes implicate differential modifier effects

Like our previous findings with age at SDMT30 and age at TMS30 landmarks (Lee et al., 2022), which fall in the earlier DCL4-TF6 interval, the relative peak heights at *MSH3* and *FAN1* were reversed in the later age at SDMTq25 and age at TMSq75 GWAS (Figure S5, SDMT and TMS, respectively), indicating a difference in the relative impact of these modifiers in cognitive and motor domains. To further explore this phenomenon, we first extended the analysis to a second UHDRS cognitive measure, Stroop Word Reading (SWR) to generate the age at SWRq25 landmark. Surprisingly, the patterns of GWAS signals detected by age at SDMTq25 and age at SWRq25 (n = 11,373, mean age 65.2 for CAG 42) revealed notable differences (Figure S5, SDMT and StroopWord, respectively). For example, unlike SDMT, the SWR landmark showed *FAN1* as the most significant locus and also detected *RRM2B* at genome-wide significance, suggesting differential contribution of these modifiers to the two cognitive phenotypes. Notably, neither cognitive landmark captured the *TCERG1* locus seen prominently by the ages at motor onset, DCL4 and TFC6 (Figure 1, Figure S1) and TMSq75 (Figure S5), further suggesting a differential influence of this modifier on the motor domain.

The composite nature of the TMS, which is the sum of 31 individual items aimed at different motor disturbances, prompted us to also test for detection of modifier loci by distinct TMS subcomponents. We defined these as in (Marder et al., 2000): bradykinesia (BRK; 11 items), chorea (CHO; seven items), dystonia (DYS; five items), oculomotor function (OCL; six items) and rigidity (RIG; two items). In GWAS, age at BRKq75 (n = 11,704; mean age 63.4 for CAG 42) showed a similar power and overall pattern to age at TMSq75 (Figure S5, bradykinesia), while age at OCLq75 (n = 11,308; mean age 66.0 for CAG 42) generally yielded less significant signals (Figure S5 oculomotor). In contrast, age at CHOq75 (n = 7,200, mean age 64.7 for CAG 42) and RIGq75 (n = 9,807, mean age 69.5 for CAG42) revealed significant evidence of modification only by *FAN1* (Figure S5, chorea and rigidity). The relatively smaller sample sizes for these motor measures reflect flattening of their mean trajectories later in the disease, such that fewer individuals are predicted to reach the q75 value observed in the overall dataset. Interestingly, age at DYSq75 (n = 10,708; mean age 59.8 for CAG 42) did not detect any significant modifier loci (Figure S5, dystonia). The paucity of significant signals across the late CHO, RIG and DYS landmarks suggests that there may be a relatively greater impact of non-genetic factors (e.g., pharmacotherapy) on the variability of these clinical phenotypes. For these and other clinical phenotypes, future genetic analyses across the entire disease trajectory, accounting where possible for non-genetic factors, considering HD individuals with longer CAG repeat lengths and comparing phenotypic extremes might provide more evidence for their genetic modifiers. Notably pathway analyses for the CHO, DYS and RIG landmark results detected mismatch repair pathways at nominal significance, but these were not the top pathways observed. However, mismatch repair pathways were at the top of the list for the late SDMT, SWR, BRK, OCL and TMS landmarks, and survived Bonferroni correction for all but age at TMSq75.

Taken together, the GWAS of the TMS subcomponent landmarks indicated that, like the cognitive measures, the different motor disturbances differentially detect and may be differentially influenced by the clinical modifiers. These landmarks are represented as z-scores calculated from distributions potentially influenced by distinct non-genetic factors, precluding direct quantitative comparison of p-values between phenotypes. However, within a single phenotype, the pattern of signals across the individual modifier effects relative to each other could be judged, allowing subsequent comparison of these relative patterns between phenotypes to implicate differential modifier impacts. We limited this comparison to the 10,793 individuals where all five informative phenotypes (SDMT, SWR, TMS, BRK and OCL) were ascertained.

Figure 6 shows the patterns of significance for the clinical modifier effects. Age at TMSq75 (red) and age at BRKq75 (blue) generally gave a similar p-value pattern, with the *FAN1* 15AM1 and 15AM2 effects being the most significant. Although the signals for the TMS landmark were typically stronger than for the BRK landmark, in a few cases (e.g., 5BM1, 19BM1, 22BM1), this order was reversed, suggesting possible differential impact relative to the other TMS subcomponents. The age at OCLq75 (gold) landmark produced a pattern of less significant p-values than the BRK and TMS phenotypes for all modifiers, with the notable exception of the new 22BM1 locus. The age at OCLq75 signal for 22BM1 was comparable to that of the TMS landmark, suggesting a proportionately higher impact of this modifier than the other loci on influencing oculomotor dysfunction, relative to modifier effects on the TMS landmark.

**Figure 6.**
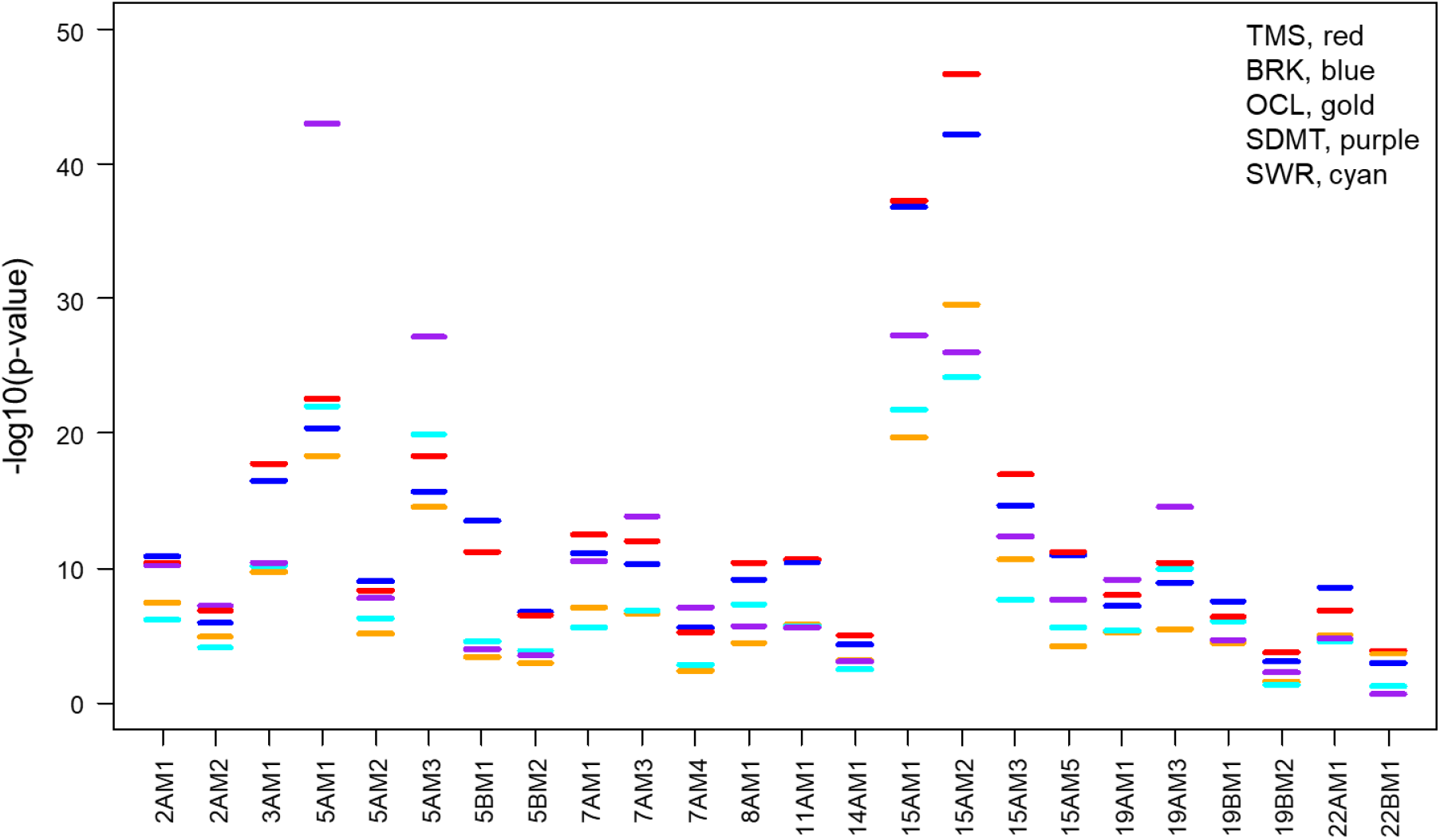
– Patterns of significance across informative late-disease landmarks. For each modifier effect detected in the clinical GWAS, the significance for the top SNV in the same set of 10,793 HD participants is plotted for each of five late HD trajectory landmarks: TMSq75 (red), BRKq75 (blue), OCLq75 (gold), SDMTq25 (purple) and SWRq25 (cyan). Comparison of the relative significance of modifier effects within and across phenotypes suggests differential impact of some modifier effects on individual cognitive and motor phenotypes.

The most striking divergence of relative signal pattern was that displayed by age at SDMTq25 (purple), which detected the frequent *MSH3* modifier 5AM1 as most significant, along with a distinct pattern for several other modifier effects. Like *MSH3* (5AM1, 5AM3), and unlike *FAN1* (15AM1, 15AM2), two of the modifier effects at *PMS2* (7AM3 and 7AM4) and at *LIG1* (19AM1 and 19AM3) showed relatively more significant signals for the SDMT than the TMS and BRK landmarks. In a contrasting pattern, at *MLH1* (3AM1), *TCERG1* (5BM1, 5BM2), *RRM2B* (8AM1), *CCDC82* (11AM1), *POLD1* (19BM1), *MED15* (22AM1) and the new 22BM1 locus, these motor landmarks yielded much more significant p-values than the SDMT landmark in the same set of individuals. Age at SWRq25 (cyan) yielded a pattern of generally weaker p-values than age at SDMTq25 except at *RRM2B* (8AM1), again suggesting a differential impact of this modifier.

Taken together with the cell-type specific regulation of CAG repeat expansion by individual modifier loci implied by their differential effects in neurons and hematopoietic stem cells, the differential patterns of modifier signals among cognitive and motor measures suggest that cell-type specific regulation of some DNA repair and other modifiers also occurs in different brain cells. The relationship between modifier effects and the phenotypes that capture them could provide the basis for identifying specific cell populations contributing to these characteristic clinical phenotypes.

## DISCUSSION

The evidence that variants in DNA-maintenance genes alter the course of HD before the onset of clinical manifestations has fueled a focus on somatic expansion of the *HTT* CAG repeat as the driver of symptom onset and the exploration of DNA repair proteins, and the processes in which they participate, as potential targets for therapeutic intervention to delay or prevent the manifestations of HD. Now GWAS of additional clinical phenotypes and of CAG expansion itself, as measured in blood DNA, support the view that a specific modifiable mismatch repair-related process, rather than a general DNA damage response, is responsible for CAG repeat expansion. Together, the clinical and molecular phenotypes detect all major mismatch repair genes as modifier loci and support *FAN1* as a suppressor of the expansion process. However, our studies also reveal an underlying complexity in terms of cell-type specificity of these somatic expansion-related modifier effects, including the differential involvement of the *MSH2/MSH6* region and the chromosome 17 locus, detected only by the somatic expansion GWAS, and genes such as *POLD1* and *LIG1,* detected only in the clinical GWAS. Moreover, new GWAS hits at *TCERG1*, *MED15,* and the 22BM1 locus, that extend the repertoire of modifiers not obviously connected to DNA repair, also imply potential cell-type specificity in the modes by which the somatically expanded CAG repeat triggers the dysfunction and loss of target neurons.

The natural and long-standing assumption that mutant huntingtin drives HD pathogenesis has led to several clinical trials to lower the expression of huntingtin. These have not yet produced a successful therapy and have been complicated individually by safety issues or lack of target engagement. However, the possibility that full-length mutant huntingtin is not the cause of neuronal dysfunction and death must be considered, so the pathogenic role, if any, of a protein product produced from the expanded *HTT* allele is of particular interest. Notably, the GWAS of HD clinical landmarks have not revealed any DNA variations tagging brain eQTLs for *HTT*. However, the ultimate driver of toxicity after critical-length somatic repeat expansion might involve the expression only of *HTT* exon 1 to produce a polypeptide dominated by polyglutamine (HTT1a) (Gipson et al., 2013; Hoschek et al., 2024). Alternatively, polyglutamine may not be involved, and repeat-associated non-AUG translation (RAN translation) to produce a different toxic polypeptide, or a mechanism involving RNA toxicity or chromatin effects may be the culprit (Bruneau and Nora, 2018; Rudich et al., 2020; Schwartz et al., 2021).

Non-canonical alleles, CAA/CCA-loss and CAACAG duplication, dramatically hasten (∼10 years) and mildly delay (∼4 years) motor onset, respectively, after adjusting for uninterrupted CAG length. Against expectation, the striking clinical effect of the CAA/CCA-loss sequence does not appear to be via a general impact on the propensity for somatic expansion of the adjacent CAG repeat. Although it would be expected by its clinical impact to be associated with increased expansion, the CAA/CCA-loss allele is instead associated with decreased CAG repeat expansion both in HD blood cell DNA and in a cell line model of CAG repeat expansion. Nor is the presence of the CAA/CCA-loss allele in HD cortex indicative of greater expansion than expected for the given CAG repeat length, though the numbers are small and an effect in a specific cell type cannot be ruled out. This strongly implies a potential impact on HD pathogenesis downstream from initial somatic CAG expansion. The non-canonical alleles could conceivably alter somatic expansion kinetics only after a critical repeat length is achieved, affect the threshold at which toxicity is triggered, or participate in the toxicity mechanism itself. Therefore, understanding the molecular basis of these *HTT* CAG 3’ non-canonical sequence effects can provide a route for distinguishing among subsequent steps in HD pathogenesis, including putative toxicity mechanisms. The same argument likely applies to the CAA-loss and CCA-loss alleles, which showed modest significance for earlier than expected motor onset despite a small numbers of carriers. Notably, the CCA-loss allele was the most frequent HD chromosome in a study of individuals of African ancestry that detected no increased CAG expansion in blood DNA and yet an earlier age at diagnosis (Dawson et al., 2022), similar to what we have observed here.

Importantly, our findings also indicate that the sequence context of the CAG repeat can be cell-specific in its effect, as illustrated by rs146151652 in the *HTT* 5’-UTR. This SNV, which shows genome-wide significant association with CAG repeat expansion of the nearby canonical CAG repeat sequence in blood DNA, alters an upstream open reading frame region suggested to regulate *HTT* mRNA translation (Lee et al., 2002). Yet, rs146151652 is not associated with motor onset or other clinical landmarks. This strongly implies that the 4ABEM1 *HTT* modifier effect influences CAG repeat expansion in hematopoietic cells, but not in neurons vulnerable to HD.

The phenomenon of cell specificity is further emphasized by genome-wide significant DNA-maintenance modifier locus variants. *MSH2/MSH6* variation is highly significant in the blood cell CAG expansion but not the clinical landmark GWAS, whereas *MLH3* is a genome-wide significant hit in the former but only suggestive significant in the latter. The converse is also observed: *MLH1, PMS1, POLD1,* and *LIG1* are hits in the clinical GWAS but are not significant in the blood somatic expansion GWAS. Since these loci are implicated in CAG repeat instability in mouse or cell models (Wheeler and Dion, 2021), the failure of some to detect significant effects on both clinical phenotypes and blood CAG expansion suggests that either the regulation of these genes or the details and stoichiometry of the CAG expansion machinery differs in HD hematopoietic stem cells and clinically relevant target neurons.

Multiple variants at *MSH3, FAN1* and *PMS2* provide a modifier allelic series at each locus, further highlighting cross-tissue similarities and differences in contributions to CAG repeat expansion. The therapeutic focus on *MSH3*, which became an early target for downregulation to reduce or block CAG repeat expansion and thereby delay or prevent HD onset of disease manifestations, appears to be bolstered by observations that the SNVs tagging the frequent *MSH3* clinical hastening effect are coincident with those associated with increased blood CAG expansion. This concordance suggests that blood CAG repeat expansion can provide a molecular biomarker for assessing potential therapeutics. On the other hand, the underlying CAG repeat expansion mechanism, which ultimately involves multiple functional partners, may be complicated, as illustrated by modifier effects at *FAN1* and *PMS2*. Although three *FAN1* modifier haplotypes that hasten clinical phenotype (including two that carry missense SNVs that alter FAN1 function) also increase CAG expansion in blood, the more common modifier effects are distinct. The top 15AM2 SNV, rs3512, in the *FAN1* 3’-UTR has been implicated in microRNA regulation of FAN1 levels (Kim et al., 2024), but this mechanism appears not to determine CAG instability in hematopoietic stem cells.

Similarly, although *PMS2* displays multiple modifier effects, only one missense SNV tags a modifier effect in both clinical and somatic expansion GWAS. The shared missense variants at *FAN1* and *PMS2* that proved consistent across the two types of GWAS suggest that structural changes can have cross-tissue influences while the other differential modifier effects likely involve cell-type specific regulation of gene expression. However, this argument does not hold for the missense variant in *MSH2*, significant only for blood CAG expansion, or for the *POLD1* missense variant and a rare missense variant in *LIG1*, both significant only for clinical modification, pointing to additional complexity in the processes underlying CAG repeat expansion. Overall, our data forecast that differences in modifier effects related to somatic expansion could ultimately reflect cell-type differences in the regulation of the modifier gene expression or activity, in the detailed operation of the mismatch repair machinery or division status, and/or in the chromatin context of the CAG repeat.

Layered onto these phenomena, whose operation in different types of HD-vulnerable neurons could similarly be the basis for relative differences in DNA-maintenance gene modifier effects across individual phenotypes (e.g., the more significant impact of *MSH3* in the cognitive domain), are the unknown interactions between genetic variants within a given cell type. Consequently, these differences could also provide a route to defining cell types in the brain that contribute differentially to particular phenotypes. This concept is not limited to the somatic expansion phase of HD pathogenesis since several modifier loci do not harbor genes known to be directly involved in DNA repair, and their potential for indirect involvement in somatic expansion is not supported by the blood CAG repeat expansion GWAS. Consequently, *TCERG1, RRM2B, CCDC82, MED15*, and the new 22BM1 locus are likely to contribute to the second phase of HD pathogenesis. Notably, these genes display the most distinct differences in relative significance across the late disease cognitive and motor landmarks, suggesting that the somatic expansion of the *HTT* CAG repeat may trigger different damaging consequences in different cell types. A more detailed examination of these modifier effects in molecular analyses that can distinguish cell types and their CAG lengths should provide a better understanding of the causes of HD pathology across the brain. On the other hand, further delineation and consideration of additional clinical phenotypes (e.g., psychiatric measures) across the entire disease trajectory from presymptomatic to end-stage HD, combined with further genetic analyses (both GWAS and comparison of extremes), will define the relative importance of modifiers in affecting the course of HD and better inform the identification and testing of potential treatments. The overall picture of HD pathogenesis as resulting from prolonged, time-dependent somatic CAG expansion and involving a complexity of cell-type specific processes leading to characteristic pathology and phenotypes is likely to apply to numerous other DNA repeat diseases whose maifestations onset in adulthood. Consequently, the path opened by shared and distinct genetic modifiers to dissect their trajectory of pathogenesis should also increase the understanding and potential for rational treatment of the growing number of other repeat disorders (Depienne and Mandel, 2021).

## ACKNOWLEDGEMENTS

Supported by CHDI Foundation, National Institutes of Health USA (NS082079, NS091161, NS016367, NS049206, NS105709, NS119471, and NS127866), the Hereditary Disease Foundation, and the Medical Research Council Centre for Neuropsychiatric Genetics and Genomics (MR/L010305/1). T.H.M. was supported by a Clinician Scientist Fellowship from the Medical Research Council, U.K. (MR/X018253/1). We thank the Harvard Tissue Bank at McLean’s Hospital, the National Neurological Research Bank at the Veteran’s Administration in Los Angeles, the New York Brain Bank at Columbia University, and the Massachusetts Alzheimer Disease Resource Center Bank at the Massachusetts General Hospital for post-mortem HD brain tissue. Biosamples and data used in this work were also generously provided by the participants in the Enroll-HD study and made available by CHDI Foundation, Inc. Enroll-HD is a clinical research platform and longitudinal observational study for Huntington’s disease families intended to accelerate progress towards therapeutics; it is sponsored by CHDI Foundation, a nonprofit biomedical research organization exclusively dedicated to collaboratively developing therapeutics for HD. Enroll-HD and the previous contributing HD studies of the Huntington Study Group, the European Huntington Disease Network and the MA HD Center Without Walls would not be possible without the vital contribution of the research participants and their families. We also thank those individuals who contributed to the collection of the Enroll-HD data, listed at https://enroll-hd.org/enrollhd_documents/ENROLL-HD_AcknowledgementsListPDS6_v1.0_20230119.pdf, and to previous HD studies, listed in the Supplementary Material of (Lee et al., 2017), and in the Supplemental Information of (GeM-HD Consortium, 2015).

## AUTHOR CONTRIBUTIONS

Conceptualization, J.F.G., M.E.M, J-M.L., V.C.W., D.G.M., L.J., P.H. and S.K.; Methodology, J.- M.L., Z.L.M., K.C., J.D.L., T.G., V.C.W., A.G., M.C., V.L., and P.H.; Resources, D.L., M.O., G.B.L., J.S.P., E.R.D., and R.H.M.,; Investigation, J-M.L., V.C.W., R.M.P., Z.L.M, K.C., J.W.S., S.L., J.- H.J., Y.L., K.-H.K., D.E.C., J.D.L., I.S.S., R.M.P., J.V.G., J.S.M., J.S., E.E., J.R., T.G., A.G., M.C., V.L., and C.W.; Data Curation, J-M.L., Z.L.M., K.C., V.C.W., R.M.P., A.G., M.C., and V.L.; Formal Analysis, J-M.L., Z.L.M., K.C., J.W.S., S.L., J-H.J., Y.L., A.G., M.C., V.L., and C.W.; Project Administration, J.F.G., M.E.M., D.G.M., L.J., and P.H.; Writing - Original Draft: J.F.G. and M.E.M.; Writing – Review and Editing: J.F.G., M.E.M., J-M.L., Z.L.M., J.D.L., V.C.W., D.G.M., A.G., M.C., V.L., C.W., T.H.M., P.H., S.K., C.S., M.O., G.B.L., J.S.P., E.R.D., and R.H.M.; Visualization, J-M.L., Z.L.M., V.C.W., A.G., M.C., and V.L.; Software: H.L., M.C., Z.L.M., J-M.L.; Supervision: J.F.G., M.E.M., J-M.L., V.C.W., D.G.M., L.J., and P.H.; Funding Acquisition: J.F.G., M.E.M, J- M.L., V.C.W., D.G.M., L.J., and P.H.

## DECLARATION OF INTERESTS

J.F.G. and V.C.W. were founding scientific advisory board members with a financial interest in Triplet Therapeutics Inc. Their financial interests were reviewed and are managed by Massachusetts General Hospital (MGH) and Mass General Brigham (MGB) in accordance with their conflict of interest policies.

J.F.G. consults for Transine Therapeutics, Inc. (dba Harness Therapeutics) and has previously provided paid consulting services to Wave Therapeutics USA Inc., Biogen Inc. and Pfizer Inc.

V.C.W. is a scientific advisory board member of LoQus23 Therapeutics Ltd. and has provided paid consulting services to Acadia Pharmaceuticals Inc., Alnylam Inc., Biogen Inc., Passage Bio and Rgenta Therapeutics. V.C.W. and R.M.P. have received research support from Pfizer Inc.

J-M.L. consults for GenKOre and serves on the scientific advisory board of GenEdit Inc. Within the last 36 months D.G.M. has been a scientific consultant and/or received an honoraria/grants from AMO Pharma, Dyne, F. Hoffman-La Roche, LoQus23, MOMA Therapeutics, Novartis, Ono Pharmaceuticals, Pfizer Pharmaceuticals, Rgenta Therapeutics, Sanofi, Sarepta Therapeutics Inc, Script Biosciences, Triplet Therapeutics, and Vertex Pharmaceuticals. D.G.M. also had research contracts with AMO Pharma and Vertex Pharmaceuticals.

J.D.L. is a paid Advisory Board member for F. Hoffmann-La Roche Ltd and uniQure biopharma B.V., and he is a paid consultant for Vaccinex Inc, Wave Life Sciences USA Inc, Genentech Inc, Triplet Inc, and PTC Therapeutics Inc.

T.H.M. is an associate member of the scientific advisory board of LoQus23 Therapeutics Ltd and has consulted for Transine Therapeutics, Inc. (dba Harness Therapeutics).

P.H. is a member of the Enroll-HD Scientific Review Committee.

S.K. and C.S. are employed by CHDI Management, Inc., as advisors to the CHDI Foundation.

E.R.D. has provided consulting services to Abbott, Abbvie, Acadia, Acorda Therapeutics, Biogen, Biohaven Pharmaceuticals, BioSensics, Boehringer Ingelheim, Caraway Therapeutics, Cerevance, CuraSen, DConsult2, Denali Therapeutics, Eli Lilly, Genentech, Health & Wellness Partners, HMP Education, Karger, KOL groups, Life Sciences Consultant, Mediflix, Medrhythms, Merck, Mitsubishi Tanabe Pharma America, Inc., MJH Holdings, NACCME Novartis, Otsuka, Praxis Medicine, Sanofi, Seelos Therapeutics, Spark Therapeutics, Springer Healthcare, Theravance Biopharmaceuticals and WebMD. He has received grant support from Averitas Pharma, Biogen, Burroughs Wellcome Fund, Pfizer, Photopharmics, and Roche and has an ownership interest in Included Health, Mediflix, SemCap and Synapticure.

G.B.L. has provided consulting services, scientific advice, scientific advisory board functions, independent data monitoring committee services and/or lectures for: Acadia Pharmaceuticals, Affiris, Allergan, Alnylam, Amarin, AOP Orphan Pharmaceuticals AG, Bayer Pharma AG, Boehringer-Ingelheim, CHDI-Foundation, Deutsche Huntington-Hilfe, Desitin, Genentech, Genzyme, GlaxoSmithKline, F. Hoffmann-LaRoche, Ipsen, ISIS Pharma (IONIS), Lilly, Lundbeck, Medesis, Medivation, Medtronic, NeuraMetrix, Neurosearch Inc., Novartis, Pfizer, Prana Biotechnology, Prilenia, PTC Therapeutics, Raptor, Remix Therapeutics, Rhône-Poulenc Rorer, Roche Pharma AG Deutschland, Sage Therapeutics, Sanofi-Aventis, Sangamo/Shire, Siena Biotech, Takeda, Temmler Pharma GmbH, Teva, Triplet Therapeutics, Trophos, UniQure and Wave Life Sciences.

R.H.M. is a paid Advisory Board member for Rgenta Therapeutics and is on the Scientific Advisory Board with financial interests in Gatehouse Bio Inc.

All other authors have nothing to declare.

## METHODS

### Human Participants

Patient consents and the overall study were reviewed and approved by the Mass General Brigham Institutional Review Board (MGB IRB). In all, we analyzed genetic data from 16,640 HD subjects. Among them, 11,261 were genotyped in previous GWA studies (GWA1-5) (GeM-HD Consortium, 2015, 2019) and 5,379 were genotyped and/or sequenced in the current study (GWA6). The data for newly analyzed subjects used in this work were generously provided by the participants in the Enroll-HD study and made available by CHDI Foundation, Inc. Enroll-HD is a global clinical research platform designed to facilitate clinical research in Huntington’s disease. Core datasets are collected annually from all research participants as part of this multi-center longitudinal observational study. Data are monitored for quality and accuracy using a risk-based monitoring approach. All sites are required to obtain and maintain local ethical approval and participants gave written informed consent. All experiments described herein were conducted in accordance with the declaration of Helsinki.

HD participant lymphoblastoid cell line DNAs used as sources of DNA for different *HTT* alleles and for MiSeq size controls were obtained from the MGB IRB-approved CHGR Neurodegenerative Repository and the Coriell Institute. SPARK whole genome sequences, available through the Simons Foundation Autism Research Initiative (SFARI) were also utilized in determining haplotypes. Human postmortem frontal cortex (BA9) tissue was obtained from the New York Brain Bank, the Harvard Brain Tissue Resource Center (Belmont, MA, USA), the National Neurological Research Bank (Los Angeles, CA, USA) and the Massachusetts Alzheimer’s Disease Research Center (ADRC) under an approved protocol by the MGB IRB.

### Cell models

hTERT-RPE1 cells (CRL-4000, ATCC, female origin) and derivatives were maintained at 37°C and 5% CO_2_ in DMEM/F-12 (Gibco), 10% fetal bovine serum (Sigma-Aldrich), and Penicillin Streptomycin (Gibco) with media changes every 2-3 days. The identity of RPE1 cells was not authenticated since we did not concurrently culture any commonly misidentified cell lines after receipt from ATCC. We previously modified RPE1 cells with a canonical *HTT* exon 1 inserted into the AAVS1 safe harbor locus to construct the RPE1-AAVS1-CAG115 system for quantifying CAG repeat instability (McLean et al., 2024). For this study, we modified hTERT-RPE1 cells with non-canonical CAACAG-dup and CAA/CCA-loss *HTT* exon 1 variants in the AAVS1 safe harbor locus (see Method Details).

### Genome-wide SNP genotyping

Six batches of GWAS data were analyzed in this study. Genotyping of GWA1-5 samples are described elsewhere (GeM 2015; GeM 2019). DNA samples for GWA6 were genotyped by Illumina Global Screen Array (https://www.illumina.com/products/by-type/microarray-kits/infinium-global-screening.html). Genotyping and genotype calling were performed at the Broad Institute using the Illumina genotyping algorithm. Since GWA data sets were genotyped using different arrays, we performed quality control (QC) analysis for each GWA independently in order to generate a dataset for subsequent genotype imputation. Briefly, we excluded SNVs that showed genotyping call rate < 95% or minor allele frequency < 1% in the typed data. We did not exclude subjects based on genetic ancestry. These data were also used to characterize the genetic ancestry using the PLINK program (https://zzz.bwh.harvard.edu/plink/) and familial relationships using the GEMMA program (https://github.com/genetics-statistics/GEMMA).

### Genotype imputation and quality control-MGH

QC-passed SNPs were further subjected to data pre-imputation QC using the “HRC or 1000G Imputation preparation and checking” program (https://www.chg.ox.ac.uk/~wrayner/tools/; Version 4.3.0). Then, genotypes were imputed separately for each GWAS1-6 set using the TOPMed Imputation Server (https://imputation.biodatacatalyst.nhlbi.nih.gov/#!; 1.5.7) involving MINIMAC4 and EAGLE V2.4. We used TOPMed r2 as the reference panel (i.e., all populations). Post-imputation, we further excluded SNVs with 1) imputation r^2^ value < 0.5 in any of the GWA datasets, 2) call rate < 100%, 3) Hardy-Weinberg equilibrium p-value < 1E-6 except for the chromosome 4:1-5,000,000 region, or 4) minor allele frequency < 0.1%. These quality control filters generated approximately 19,000,000 imputed SNVs.

### Genotype imputation and quality control-Glasgow

Due to restrictions on sharing certain individual data between groups, imputation on GWA6 was performed separately in Glasgow. For the pre-imputation quality control, samples with > 5% of missing genotypes, and SNVs with > 5% of missing genotypes or minor allele frequency of < 1% were filtered out using plink1.9. The genotype file was converted to vcf files using bcftools. The vcf files were uploaded to TOPMed and imputation was run on unphased input to GRCh37/hg19. The vcf files were downloaded post imputation and merged together using bcftools. The merged file was filtered to exclude SNVs with imputation quality score (RSq) < 0.5 using bcftools. The vcf file was converted to a genotype file in binary plink format using plink1.9. The genotype file was filtered to exclude variants with minor allele frequency < 0.1%, Hardy-Weinberg equilibrium p-value < 1.0E-06 and genotype call rate <100%. These quality control filters generate ∼15,700,000 imputed SNVs.

### DNA extraction from postmortem brain

Approximately 25 μg of flash frozen BA9 cortical tissue was rapidly chopped into smaller pieces and genomic DNA was extracted using the Qiagen DNeasy Blood & Tissue kit, according to the manufacturer’s spin-column protocol. Briefly, finely chopped tissue was placed into a 1.5 ml microcentrifuge tube, then 20 µl Proteinase K reagent was added and mixed by vortexing, then incubated at 56°C for about 1-3 hours until lysis was complete after which the sample was vortexed for 15 seconds and 200 µl Buffer AL was added, followed by vortexing to mix, followed by the addition of 200 µl ethanol (96–100%), and mixing. This was then pipetted onto a DNeasy Mini spin column that had been placed in the 2 ml collection tube, followed by centrifugation at ≥ 6000 x g for 1 minute. The DNeasy Mini spin column was then placed in a fresh 2 ml collection tube and column washed by adding 500 µl Buffer AW1, followed by centrifugation for 1 min at ≥ 6000 x g. The DNeasy Mini spin column was again placed in a new 2 ml collection tube and 500 µl Buffer AW2 was added, followed by centrifugation for 3 minutes at 20,000 x g (14,000 rpm). DNA was then eluted from the dry column membrane by putting the DNeasy spin column in a fresh 1.5 ml microcentrifuge tube and pipetting 200 µl Buffer AE directly onto the DNeasy membrane, held at room temperature for 1 minute and eluant collected by centrifugation for 1 minute at ≥ 6000 x g (8000 rpm). DNA prepared in this manner was then utilized directly for genotyping and CAG expansion ratio assays.

### MGH HTT ABI and MiSeq CAG genotyping

For all samples utilized in this study, the PCR amplification assay reported by Warner et. al., 1993, modified for the ABI3730XL DNA sequencer, was used to estimate *HTT* CAG repeat size, relative to sequenced DNA standards of known uninterrupted (‘pure’) CAG repeat lengths. Reactions: 10 µl with 1.25 mM MgCl_2_ (Applied Biosystems); 1X Buffer II (Applied Biosystems); 0.05 U Amplitaq Gold (Applied Biosystems); 0.25 mM dNTPs (GE Healthcare); 1.2 µl of DMSO (Sigma); 0.125 µM each primer; and 80 ng of genomic DNA. The forward primer was labeled with 6FAM (HD-1) 5’ ATGAAGGCCTTCGAGTCCCTCAAGTCCTTC 3’ and the reverse primer was tailed (HD-3) 5’ GGCGGTGGCGGCTGTTGCTGCTGCTGCTGC 3’. Conditions were: 94°C for 4 minutes, then 35 X denaturation (94°C 30 seconds), annealing (65°C 30 seconds), extension (72°C 45 seconds), and final extension (72°C 10 minutes), holding at 15°C. Products were resolved on an ABI 3730XL DNA Analyzer (36 cm array, POP-7 Polymer, standard fragment analysis conditions), together with internal size standard ladder where 0.8 µl PCR product was loaded in 9.4 µl Hi-Di Formamide (Applied Biosystems), with 0.1 µl GeneScan 500 LIZ (Applied Biosystems). The .fsa files generated were analyzed with GeneMapper v5.0 (Applied Biosystems) software. The *HTT* CAG repeat allele sizes for samples were estimated relative to the fragment sizes of the sequenced *HTT* CAG repeat genomic DNA standards, based upon the highest peak-signal (main peak) in the normal and the expanded CAG repeat range in each trace.

For the clinical GWAS, the pure CAG size and configuration of adjacent sequence, to identify canonical and non-canonical alleles, was determined from MiSeq data generated at MGH and Q^2^ Solutions/EA Genomics (Durham, NC, USA). The generation of the latter data is described in detail in the next section. MGH analyzed both sets of data for the clinical GWAS while the Glasgow group separately analyzed the Q^2^ Solutions/EA Genomics data for the MiSeq somatic expansion GWAS, as described in the next section.

MiSeq performed at MGH utilized 2 x 300bp Illumina MiSeq Adapted 16S Metagenomic Targeted Resequencing Protocol Library Preparation guide (Part # 15044223 Rev. B). The specific primer set utilized in Step 1: oligonucleotide pair (PCR 1) comprising the forward and reverse *HTT* CAG target sequences and Illumina Adapter sequence (in italic text) was: ms_hd_f 5’ *TCGTCGGCAGCGTCAGATGTGTATAAGAGACAG*ATGAAGGCCTTCGAGTCCC 3’ ms_hd_r 5’ *GTCTCGTGGGCTCGGAGATGTGTATAAGAGACAG*GGCTGAGGAAGCTGAGGA 3’.

The reaction volume was 20 µl: 1X PCR Buffer (Qiagen); 1X Q-Solution (Qiagen), 200 µM dNTP (Qiagen); 0.25 µM each forward and reverse primer, 0.5 units of Taq DNA Polymerase (Qiagen); with 80 ng of genomic DNA. Cycles were: initial denaturation (93°C 3 minutes), 33X denaturation (93°C 30 seconds), annealing (60°C 30 seconds), extension (68°C 90 seconds) with final extension (72°C 10 minutes) and hold (15°C until bead clean-up). The Step 2 PCR reaction, incorporating Nextera XT Index Kit v2 index kits A, B,C and D, was in 50 ul:1X PCR Buffer (Qiagen), 1X Q-Solution (Qiagen), 200 µM dNTP (Qiagen), 0.8 µM each forward and reverse primer, 1.25 units of Taq DNA Polymerase (Qiagen), and 5 µl of Step 1 PCR product. Cycling was initial denaturation (93°C 3 minutes), 8X denaturation (93°C 30 seconds), annealing (60°C for 30 seconds), extension (68°C 90 seconds) and hold (15°C until bead clean-up). Bead clean-up was done using AMPure XP bead based reagent (Beckman Coulter), after each of the two PCR cycles with a 1X bead to PCR product ratio followed by two 80% ethanol washes. Libraries were eluted from beads with 10 mM Tris-HCl pH8.5. After clean-up, QC on each library was performed by electrophoresis on an Agilent 4200 Tape Station System (Agilent) to confirm the expected PCR amplification peaks and lack of contamination and adaptor dimer peaks. The QC-qualified libraries were then pooled, in sets of 384, re-run on the Agilent 4200 Tape Station to confirm peaks and determine concentration.

Pooled libraries were then diluted and run on the Illumina MiSeq Sequencing System (Illumina), according to Illumina guidelines, at concentrations 8 pM with 15% PhiX spike-in for a 2 x 300 base read.

Analysis to determine pure CAG size and adjacent sequence was as follows. The 300 bp MiSeq data (FASTQ format) were utilized for analysis to determine the size of uninterrupted CAG repeat and adjacent DNA sequence, by processing each sequence read pair using Python (v3.9). The forward strand start was the CAGCAGCAG 9mer (5’ end of the uninterrupted CAG repeat tract), continuing to the CAGCTTCCT 9mer (3’ end of the polymorphic triplet repeat tract) or until the end of the read and the reverse complement of the forward strand reads were analyzed in the same manner. Paired reads in which the uninterrupted CAG repeat tract and adjacent sequence matched were aggregated and counted, discarding unmatched reads. The CAG repeat genotype for each sample was generated from the uninterrupted CAG repeat allele assigned (Python v3.9), using the distribution of uninterrupted CAG repeat sizes, by identifying the two most frequent CAG peaks, Then the highest frequency adjacent sequences that contained each uninterrupted CAG repeat allele were used to create a complete genotype for both alleles of each sample, with an assigned pure CAG tract and adjacent sequence. Samples failed QC if the total paired read count was <1,000 or if paired read count for either the normal range or expanded range CAG was <200. For each sample, the CAG sizes were cross-checked against the ABI assay genotype, both to correct non-canonical allele sizes (evident from the MiSeq adjacent sequence genotype) misestimated in the ABI assay and to determine if the mismatch reflected a failure to pass QC or a primer polymorphism preventing amplification of an allele, in one or the other assay. For the clinical GWAS, the uninterrupted pure CAG size from MiSeq was utilized, except in samples for which MiSeq genotype was not available, in which case the ABI fragment assay allele sizes were utilized.

### Q^2^ Solutions MiSeq and Glasgow CAG analysis

At Q^2^ Solutions/EA Genomics, *HTT* exon 1 repeat region MiSeq was performed using the amplicon sequencing protocol described in Ciosi et al. (2018), including MiSeq library preparation, PCR, clean-up and QC, for blood DNA of 5,876 Enroll-HD participants using MiSeq-compatible PCR primers incorporating Nextera XT Index Kit v2 indexes (384 different amplicons per MiSeq run 2 x 300 base read). This protocol utilizes 40 different oligonucleotide primers containing full-length Illumina indexed adapters (in italicized text with Nextera XT Index in bold), heterogeneity spacer (in unformatted text in smaller letters) and *HTT* locus-specific primer sequence (in underlined text). These are:

MiSeq F HD319F S502: AATGATACGGCGACCACCGAGATCTACAC**CTCTCTAT**ACACTCTTTCCCTACACGACGCTC TTCCGATCTAATATAT<u>GCGACCCTGGAAAAGCTGATGA</u>

MiSeq F HD319F S503: AATGATACGGCGACCACCGAGATCTACAC**TATCCTCT**ACACTCTTTCCCTACACGACGCTC TTCCGATCTATACTT<u>GCGACCCTGGAAAAGCTGATGA</u>

MiSeq F HD319F S505: AATGATACGGCGACCACCGAGATCTACAC**GTAAGGAG**ACACTCTTTCCCTACACGACGCT CTTCCGATCTCT<u>GCGACCCTGGAAAAGCTGATGA</u>

MiSeq F HD319F S506: AATGATACGGCGACCACCGAGATCTACAC**ACTGCATA**ACACTCTTTCCCTACACGACGCTC TTCCGATCTTCC<u>GCGACCCTGGAAAAGCTGATGA</u>

MiSeq F HD319F S507: AATGATACGGCGACCACCGAGATCTACAC**AAGGAGTA**ACACTCTTTCCCTACACGACGCT CTTCCGATCTCGAT<u>GCGACCCTGGAAAAGCTGATGA</u>

MiSeq F HD319F S508: AATGATACGGCGACCACCGAGATCTACAC**CTAAGCCT**ACACTCTTTCCCTACACGACGCTC TTCCGATCTG<u>GCGACCCTGGAAAAGCTGATGA</u>

MiSeq F HD319F S510: AATGATACGGCGACCACCGAGATCTACAC**CGTCTAAT**ACACTCTTTCCCTACACGACGCTC TTCCGATCTTATTA<u>GCGACCCTGGAAAAGCTGATGA</u>

MiSeq F HD319F S511: AATGATACGGCGACCACCGAGATCTACAC**TCTCTCCG**ACACTCTTTCCCTACACGACGCTC TTCCGATCT<u>GCGACCCTGGAAAAGCTGATGA</u>

MiSeq F HD319F S513: AATGATACGGCGACCACCGAGATCTACAC**TCGACTAG**ACACTCTTTCCCTACACGACGCTC TTCCGATCT<u>GCGACCCTGGAAAAGCTGATGA</u>

MiSeq F HD319F S515: AATGATACGGCGACCACCGAGATCTACAC**TTCTAGCT**ACACTCTTTCCCTACACGACGCTC TTCCGATCTGCTT<u>GCGACCCTGGAAAAGCTGATGA</u>

MiSeq F HD319F S516: AATGATACGGCGACCACCGAGATCTACAC**CCTAGAGT**ACACTCTTTCCCTACACGACGCTC TTCCGATCTC<u>GCGACCCTGGAAAAGCTGATGA</u>

MiSeq F HD319F S517: AATGATACGGCGACCACCGAGATCTACAC**GCGTAAGA**ACACTCTTTCCCTACACGACGCT CTTCCGATCTTG<u>GCGACCCTGGAAAAGCTGATGA</u>

MiSeq F HD319F S518: AATGATACGGCGACCACCGAGATCTACAC**CTATTAAG**ACACTCTTTCCCTACACGACGCTC TTCCGATCTTAATATT<u>GCGACCCTGGAAAAGCTGATGA</u>

MiSeq F HD319F S520: AATGATACGGCGACCACCGAGATCTACAC**AAGGCTAT**ACACTCTTTCCCTACACGACGCTC TTCCGATCTCATATA<u>GCGACCCTGGAAAAGCTGATGA</u>

MiSeq F HD319F S521: AATGATACGGCGACCACCGAGATCTACAC**GAGCCTTA**ACACTCTTTCCCTACACGACGCTC TTCCGATCTATC<u>GCGACCCTGGAAAAGCTGATGA</u>

MiSeq F HD319F S522: AATGATACGGCGACCACCGAGATCTACAC**TTATGCGA**ACACTCTTTCCCTACACGACGCTC TTCCGATCTATACT<u>GCGACCCTGGAAAAGCTGATGA</u>

MiSeq R 33935.5 N701: CAAGCAGAAGACGGCATACGAGAT**TCGCCTTA**GTGACTGGAGTTCAGACGTGTGCTCTTC CGATCTA<u>AGCAGCGGCTGTGCCTGC</u>

MiSeq R 33935.5 N702: CAAGCAGAAGACGGCATACGAGAT**CTAGTACG**GTGACTGGAGTTCAGACGTGTGCTCTTC CGATCAT<u>AGCAGCGGCTGTGCCTGC</u>

MiSeq R 33935.5 N703: CAAGCAGAAGACGGCATACGAGAT**TTCTGCCT**GTGACTGGAGTTCAGACGTGTGCTCTTC CGATCTA<u>AGCAGCGGCTGTGCCTGC</u>

MiSeq R 33935.5 N704: CAAGCAGAAGACGGCATACGAGAT**GCTCAGGA**GTGACTGGAGTTCAGACGTGTGCTCTTC CGATCG<u>AGCAGCGGCTGTGCCTGC</u>

MiSeq R 33935.5 N705: CAAGCAGAAGACGGCATACGAGAT**AGGAGTCC**GTGACTGGAGTTCAGACGTGTGCTCTTC CGATCG<u>AGCAGCGGCTGTGCCTGC</u>

MiSeq R 33935.5 N706: CAAGCAGAAGACGGCATACGAGAT**CATGCCTA**GTGACTGGAGTTCAGACGTGTGCTCTTC CGATCAT<u>AGCAGCGGCTGTGCCTGC</u>

MiSeq R 33935.5 N707: CAAGCAGAAGACGGCATACGAGAT**GTAGAGAG**GTGACTGGAGTTCAGACGTGTGCTCTTC CGATCTA<u>AGCAGCGGCTGTGCCTGC</u>

MiSeq R 33935.5 N710: CAAGCAGAAGACGGCATACGAGAT**CAGCCTCG**GTGACTGGAGTTCAGACGTGTGCTCTTC CGATC<u>AGCAGCGGCTGTGCCTGC</u>

MiSeq R 33935.5 N711: CAAGCAGAAGACGGCATACGAGAT**TGCCTCTT**GTGACTGGAGTTCAGACGTGTGCTCTTC CGATCAT<u>AGCAGCGGCTGTGCCTGC</u>

MiSeq R 33935.5 N712: CAAGCAGAAGACGGCATACGAGAT**TCCTCTAC**GTGACTGGAGTTCAGACGTGTGCTCTTC CGATCTA<u>AGCAGCGGCTGTGCCTGC</u>

MiSeq R 33935.5 N714: CAAGCAGAAGACGGCATACGAGAT**TCATGAGC**GTGACTGGAGTTCAGACGTGTGCTCTTC CGATCAT<u>AGCAGCGGCTGTGCCTGC</u>

MiSeq R 33935.5 N715: CAAGCAGAAGACGGCATACGAGAT**CCTGAGAT**GTGACTGGAGTTCAGACGTGTGCTCTTC CGATCTA<u>AGCAGCGGCTGTGCCTGC</u>

MiSeq R 33935.5 N716: CAAGCAGAAGACGGCATACGAGAT**TAGCGAGT**GTGACTGGAGTTCAGACGTGTGCTCTTC CGATCAT<u>AGCAGCGGCTGTGCCTGC</u>

MiSeq R 33935.5 N718: CAAGCAGAAGACGGCATACGAGAT**GTAGCTCC**GTGACTGGAGTTCAGACGTGTGCTCTTC CGATCG<u>AGCAGCGGCTGTGCCTGC</u>

MiSeq R 33935.5 N719: CAAGCAGAAGACGGCATACGAGAT**TACTACGC**GTGACTGGAGTTCAGACGTGTGCTCTTC CGATCTA<u>AGCAGCGGCTGTGCCTGC</u>

MiSeq R 33935.5 N720: CAAGCAGAAGACGGCATACGAGAT**AGGCTCCG**GTGACTGGAGTTCAGACGTGTGCTCTTC CGATC<u>AGCAGCGGCTGTGCCTGC</u>

MiSeq R 33935.5 N721: CAAGCAGAAGACGGCATACGAGAT**GCAGCGTA**GTGACTGGAGTTCAGACGTGTGCTCTTC CGATCG<u>AGCAGCGGCTGTGCCTGC</u>

MiSeq R 33935.5 N722: CAAGCAGAAGACGGCATACGAGAT**CTGCGCAT**GTGACTGGAGTTCAGACGTGTGCTCTTC CGATCG<u>AGCAGCGGCTGTGCCTGC</u>

MiSeq R 33935.5 N723: CAAGCAGAAGACGGCATACGAGAT**GAGCGCTA**GTGACTGGAGTTCAGACGTGTGCTCTTC CGATCG<u>AGCAGCGGCTGTGCCTGC</u>

MiSeq R 33935.5 N724: CAAGCAGAAGACGGCATACGAGAT**CGCTCAGT**GTGACTGGAGTTCAGACGTGTGCTCTTC CGATCG<u>AGCAGCGGCTGTGCCTGC</u>

MiSeq R 33935.5 N726: CAAGCAGAAGACGGCATACGAGAT**GTCTTAGG**GTGACTGGAGTTCAGACGTGTGCTCTTC CGATCAT<u>AGCAGCGGCTGTGCCTGC</u>

MiSeq R 33935.5 N727: CAAGCAGAAGACGGCATACGAGAT**ACTGATCG**GTGACTGGAGTTCAGACGTGTGCTCTTC CGATCTA<u>AGCAGCGGCTGTGCCTGC</u>

MiSeq R 33935.5 N728: CAAGCAGAAGACGGCATACGAGAT**TAGCTGCA**GTGACTGGAGTTCAGACGTGTGCTCTTC CGATCAT<u>AGCAGCGGCTGTGCCTGC</u>

MiSeq R 33935.5 N729: CAAGCAGAAGACGGCATACGAGAT**GACGTCGA**GTGACTGGAGTTCAGACGTGTGCTCTTC CGATCG<u>AGCAGCGGCTGTGCCTGC</u>

PCR was performed in 384-well plate format, with a CAG-expansion canonical repeat DNA (NA13503 Coriell Institute) and a non-expanded non-canonical *HTT* repeat control DNA (NA20342 Coriell Institute) sample and two no-template control on each plate, as follows: 15 µl reaction volume contained 20 ng of blood DNA, 0.666 µM of each forward (HD319F) and reverse (33935.5) MiSeq-compatible PCR primers (using a unique index pair for each sample or control), 10% (v/v) DMSO, 1X ‘Custom PCR Master Mix, 0.048% (v/v) βME’ and 1 U of Taq DNA polymerase. PCR cycling conditions were: (96°C, 5 minutes), 28 cycles of (96°C, 45 seconds), (58.5°C, 45 seconds) and (72°C, 3 minutes), (72°C, 10 minutes). DNA concentrations associated with the following post-PCR QC steps were estimated on a Qubit fluorometer using the Qubit® dsDNA HS Assay Kit. QC was performed on 1 µl of control PCR products (0.5 ng/µl dilution) to confirm PCR amplification and absence of PCR contamination, by Bioanalyzer capillary electrophoresis. Clean-up on a pool of the 384 PCR products (5 µl each) was done in two rounds of AMPure XP Bead-Based Reagent clean-up (0.6X and 1.4X AMPure XP Bead-Based Reagent/PCR product pool ratio respectively). A 0.5 ng/µl (Qubit fluorometer using the Qubit® dsDNA HS Assay Kit) aliquot of the purified sequencing library was then run on Bioanalyzer to check expected library size and average library size. The molarity of each purified library was then determined based on the Bioanalyzer-estimate of the average fragment size in the library and the quantity of sequenceable molecules estimated by qPCR using the KAPA Library Quantification Kit. Sequencing followed Illumina guidelines, using the MiSeq Reagent Kit v3 to produce 2X300 bases per cluster with a cluster density of 1000k cluster mm^-2^ but with 5% PhiX spike-in.

Glasgow analysis of the Q^2^ Solutions/EA Genomics MiSeq reads used the alignment-based approach in ScaleHD (version 1 - https://pypi.org/project/ScaleHD in combination with cutadapt version 1.18 and bwa version 0.7.17-r1188), as described previously (Ciosi et al., 2019); (see below configuration file used) and the alignment-free approach implemented in RGT (https://github.com/hossam26644/RGT; see below the RGT settings file used on the forward/R1 MiSeq reads). RGT extracts repeat sequences from between the user-defined upstream and downstream flanking sequences, and then a 3 bp sliding window is used to count repeat units per read while allowing for sequence interruptions. RGT records the abundance of different repeat units in the sample and the identified abundance peaks are used to distinguish between germline modal variants and variants resulting from somatic mosaicism or PCR slippage. RGT provides plots to enable visual review of the peak profiles.

ScaleHD HTT genotyping configuration file:

<config data_dir=“Miseq_data” forward_reference=“4k-HD-INTER.fa” reverse_reference=“4k-HD-Reverse.fasta”>

<instance_flags demultiplex=“True” quality_control=“True” sequence_alignment=“True” atypical_realignment=“True” genotype_prediction=“True” snp_calling=“True”/>

<demultiplex_flags forward_adapter=“GCGACCCTGG” forward_position=“5P” reverse_adapter=“GCAGCGGCTG” reverse_position=“5P” error_rate=“0” min_overlap=“10” min_length=“” max_length=“”/>

<trim_flags trim_type=“Adapter” quality_threshold=“5” adapter_flag=“-a” forward_adapter=“GATCGGAAGAGCACACGTCTGAACTCCAGTCAC” reverse_adapter=“AGATCGGAAGAGCGTCGTGTAGGGAAAGAGTGT”

error_tolerance=“0.39”/>

<alignment_flags min_seed_length=“19” band_width=“100” seed_length_extension=“1.5” skip_seed_with_occurrence=“500” chain_drop=“0.50” seeded_chain_drop=“0” seq_match_score=“1” mismatch_penalty=“4” indel_penalty=“6,6” gap_extend_penalty=“4,4” prime_clipping_penalty=“5,5” unpaired_pairing_penalty=“17”/>

<prediction_flags snp_observation_threshold=“2” quality_cutoff=“1000”/>

RGT settings file:

{

“start_flank”: “GAGTCCCTCAAGTCCTTC”, “end_flank”: “CTTCCTCAGCCGCCG”,

“repeat_units”: [“CAG”,“CAA”, “CCG”, “CCA”, “CCT”],

“unique_repeat_units”:[“CAG”],

“grouping_repeat_units”:[“CAACAG”, “CCGCCA”,“CAG”, “CCG”, “CCT”], “min_size_repeate”: 5,

“max_interrupt_tract”: 5, “discard_reads_with_no_end_flank”: “True”, “discarded_reads_flag_percentage”: 10, “reverse_strand”:“False”, “minimum_no_of_reads”: 30, “3D_plot_parameters”:{

“x_units”:[“CAG”],

“z_units”:[“CCG”],

“before_x_seq”:“”, “after_x_seq”:“CAACAG”, “before_z_seq”:“”, “after_z_seq”:“CCT”

}

Alignment data were manually checked if: software genotypes did not agree with each other; ScaleHD failed to genotype; no CAG-expanded allele was detected or if the allele was > 48 CAGs; the % of aligned forward reads was < 90%; or a novel non-canonical allele was detected. The RGT 3D graphical output of the repeat length distributions (x-axis = number of CAGs, y-axis = number of CCGs and z-axis = number of reads) was used to visually inspect the repeat length frequency distributions for all samples to identify unexpected patterns (*e.g.,* distributions with more than two peaks/modes). At that stage, MiSeq data was excluded for 893 participants because the MiSeq data presented > two alleles (13 participants’ samples), less reads than the no-template controls (one participant’s sample), because genotyping from MiSeq data had failed (472 participants’ samples), because the CAG lengths obtained from the MiSeq data were not concordant with the CAG lengths available from the Enroll-HD study (85 participants’ samples) or because the sequencing quality in one of the MiSeq runs did not meet Illumina standards (an additional 322 participants’ samples).

### Generation of cell models for CAG expansion

We generated hTERT-RPE1 derivative lines containing non-canonical CAACAG-dup and CAA/CCA-loss *HTT* exon 1 variants at the AAVS1 locus to match the RPE1-AAVS1-CAG115 cell line with a canonical *HTT* exon 1 sequence (McLean et al., 2024). Exon 1 fragments were derived from HD participant lymphoblastoid cells and expanded in plasmids to matching uninterrupted CAG repeat lengths. The entire *HTT* exon 1 coding sequence was amplified using UltraRun® LongRange PCR Kit (QIAGEN, 206442) with 10% DMSO using the following PCR conditions: denaturation 93°C (3 min), 35 cycles of 93°C (30 s), 61°C (15 s), 68°C (60 s), with a final extension of 72 °C (10 min), using primers with SalI cloning sites (F: CATGTACGgtcgacaccgccATGGCGACCCTGGAAAAGCTG, R: CATGTACGgtcgacTCGGTGCAGCGGCTCCTC). Amplicons were cloned into a plasmid for targeting the AAVS1 safe harbor locus (AAVS1-TRE3G-EGFP was a gift from Su-Chun Zhang (Addgene plasmid # 52343; http://n2t.net/addgene:52343; RRID:Addgene_52343)). Plasmids carrying CAACAG-dup and CAA/CCA-loss *HTT* exon 1 variants were expanded from ∼40 to 115 uninterrupted repeats by precisely cutting the plasmid at the CAG repeat sequence with CRISPR-Cas9 and recombining with Gibson assembly. A plasmid with 115 uninterrupted CAG repeats and a canonical sequence at the 3’ end of the repeat was used to isolate a fragment with 115 CAG repeats. CRISPR-Cas9 crRNA sequence (GGCGGTGGCGGCTGTTGCTG, Integrated DNA Technologies), was duplexed with tracrRNA (Integrated DNA Technologies) and a ribonucleoprotein (RNP) complex made by combining 10 μM duplex with 10 μM Cas9 (Integrated DNA Technologies) and incubated at 25°C for 10 min. A total of 10 μL RNP complex was combined with 2 μg plasmid DNA, SpeI restriction enzyme (for second Gibson homology site, New England Biolabs), and CutSmart buffer (New England Biolabs) up to 30 μL and digested overnight. The same approach was taken for plasmids with CAACAG-dup and CAA/CCA-loss *HTT* exon 1 variants, but the CAG repeat was targeted with crRNA sequence (CTGCTGCTGCTGCTGCTG, Integrated DNA Technologies) and PspXI restriction enzyme (New England Biolabs). The 0.8 kb fragment with 115 uninterrupted repeats and 12 kb digested fragment with the CAACAG-dup and CAA/CCA-loss variants contain homology sequences within the CAG repeat itself (3’ and 5’ ends of fragments, respectively) and a 20 bp sequence of plasmid backbone. NEBuilder® HiFi DNA Assembly Master Mix (New England Biolabs) was used following the manufacturer’s protocol to recombine the fragments. The resulting plasmids with matching uninterrupted CAG repeat sequences were nanopore sequenced (Plasmidsaurus, SNPsaurus LLC) to confirm sequence identity and the repeat size was quantified by fragment analysis as described in the *HTT* CAG repeat genotyping section.

The plasmids containing different *HTT* exon 1 variants were knocked into the AAVS1 site using the standard safe harbor targeting approach. We used nucleofection (4D-Nucleofector X Unit and P3 4D-Nucleofector™ X Solution, Lonza) to transfect both the AAVS1 targeting plasmids described above and transcription activator-like effector nucleases (hAAVS1 TALEN Left and Right were gifts from Su-Chun Zhang, Addgene plasmid # 52341 & 52342; http://n2t.net/addgene:52341; http://n2t.net/addgene:52342; RRID:Addgene_52341; RRID:Addgene_52342). After 20 µg/mL puromycin selection for 1 week, clones were isolated by limiting dilution and screened for targeted integration into the locus with PCR assays on the 5’ integration site (F: CTGCCGTCTCTCTCCTGAGT, R: GTGGGCTTGTACTCGGTCAT), 3’ site (F: TGAAGCCCCTTGAGCATCTG, R: CCTGGGATACCCCGAAGAGT), and a PCR spanning the integration site to confirm mono-allelic integration (F: CTGCCGTCTCTCTCCTGAGT, R: AGGAAGGAGGAGGCCTAAGG). We used GoTaq G2 (Promega) with denaturation 94°C (2 min), 35 cycles of 94°C (30 s), 60°C (30 s), 72°C (60 s), with a final extension of 72°C (5 min), with 10% DMSO supplementation for 5’ assay and 1 M betaine (Sigma-Aldrich) for the spanning assay.

### Cell line CAG expansion assay

CAG repeat instability was compared for cell clones with canonical, CAACAG-dup and CAA/CCA-loss *HTT* exon 1 insertions in the AAVS1 locus. Cells were seeded onto a 96-well plate at high density (50,000 per well) to ensure that the instability experiment was performed in confluent non-dividing cultures. A sample was taken at day-zero for each clone with three wells per clone maintained for four weeks. DNA was isolated using Quick-DNA 96 Kit (Zymo Research) and the repeat sequence amplified in a nested PCR approach to specifically amplify the transgene and avoid competition of the normal repeat length at the endogenous *HTT* locus. The first PCR amplified from *HTT* to *GFP* (F: ATGAAGGCCTTCGAGTCCCTCAAGTCCTTC, R: GTCCAGCTCGACCAGGATG) Taq PCR Core Kit with Q solution (Qiagen) with 5 µL of the genomic DNA with initial denaturation 95°C (5 min), 12 cycles of 95°C (30 s), 65°C (30 s), 72°C (1 min 30 s), final extension 72°C (10 min). The second standard repeat sizing PCR (F: /56-FAM/ATGAAGGCCTTCGAGTCCCTCAAGTCCTTC, R: GGCGGCTGAGGAAGCTGAGGA) used 2 μl of the first as a template and had 22 cycles. The amplicons were analyzed by capillary electrophoresis using 3730XL DNA Analyzer as described above and GeneMapper 5.0 was used to assign bp size and identify fragment peaks.

### Clinical GWAS

HD age at motor onset was the age at which otherwise unexplained extrapyramidal motor signs emerged as assessed by an expert rater in the Unified Huntington’s Disease Rating Scale (UHDRS). In the rare cases where no expert rater estimate was available, the estimate was provided by family members of the participant. For the age at motor onset GWAS phenotype we calculated a CAG-specific z-score based on the median and standard deviation of the age at motor onset distribution. Sex was not significantly associated with age at motor onset and therefore, it was not included when we calculated the z-score of motor onset.

In contrast, z-scores of age at TFC6, age at DCL4 and age at clinical measure quantiles were calculated separately for males and females to correct for the impact of sex. For age at TFC6 and age at DCL4, the algorithmic predictions were those generated in a previous study using both survival analysis and longitudinal analysis to integrate clinical measures from HD natural history studies (Lee et al., 2022). Briefly, a parametric survival curve was generated for each combination of CAG/sex/education (low= high school or less, high= more than high school) for each recorded CAG length from 40 to 50 to determine the age at median survival for reaching TFC6 and for reaching DCL4. These ages were then positioned on a linear mixed model (LMM) estimate of the mean trajectory of the cUHDRS score designed for monitoring disease progression by combining UHDRS measures (TFC, TMS, SDMT and SWR). The comparison of the median survival ages for TFC6 and DCL4 to this trajectory identified a corresponding cUHDRS score for each clinical landmark. When individual trajectories of cUHDRS were estimated for each participant and extrapolated beyond their period of clinical data collection, the ages at which each individual curve achieved these TFC6 and DCL4 cUHDRS scores were designated as the age at TFC6 and age at DCL4 for that individual. We also generated landmarks based on individual cognitive measures (SDMT, SWR), TMS and subcomponents of TMS (BRK, CHO, DYS, OCL, RIG). For SDMT and SWR, which decrease with HD progression, we calculated the 25% quantile of all scores observed in the natural history dataset used to predict age at TFC6 and age at DCL4. For TMS and its subcomponents, which increase with HD deterioration, we determined the 75% quantile. We then predicted the age at each landmark for each individual as above for age at TFC6 and age at DCL4, except that the LMM estimated trajectory for the particular measure was used rather than that for cUHDRS. For the GWAS phenotype, we converted each prediction to a Z-score using the median and standard deviation of the ages for all the participants in the same CAG/sex/education group (thereby adjusting for these variables).

Uninterrupted CAG repeat lengths for the clinical GWAS were determined by MiSeq at MGH. For age at motor onset, we analyzed HD participants with 40-55 CAGs while for algorithmically predicted landmarks we analyzed HD participants with 40-50 CAGs. For each age landmark phenotype, we analyzed all individuals, regardless of ancestry. For each test phenotype, we performed combined association analysis in a mixed effect model. Briefly, the phenotype was modeled as a function of minor allele count of a test SNP, sex, batch of GWAS, and the first four principal component values (from the genetic ancestry analysis) in a linear mixed effect model with relationship matrix using the GEMMA program (version, 0.94 beta; http://www.xzlab.org/software.html) (Zhou and Stephens, 2012). Since eigenvalues based on principal components analysis appeared to level off starting at the third principal component in the scree plot analysis of GWA1-5 data (Lee et al., 2022), four principal component values were considered sufficient to correct for effects of population stratification. To avoid spurious associations due to phenotypic outliers, the main GWAS analysis was limited to SNVs with MAF > 1%, except at established loci with previous robust signals for SNVs between 0.1% and 1%: *HTT*, *MSH3*, *LIG1* and *FAN1*.

### ABI trace blood CAG PPS phenotype

For quantification of somatic CAG repeat expansion, analysis of peaks in traces for 11,377 Enroll-HD and 774 Registry individuals was performed on ABIF format FSA files produced by ABI DNA Sequence Analyzer using software libraries for Python (v3.x). The BioPython software tools (v1.7x) were used for opening and parsing the ABIF files and extracting out trace and ladder signals. Numpy (v1.19+) and scipy (v1.2+) and pandas (v0.24+) were used for ladder analysis and linear approximation for base pair sizing. Peak detection of the trace of the fragment analysis was done by using scipy (v1.2+) to perform a fast Fourier transform (FFT) on the main signal to filter primary fragment peaks around the main allelic peaks of each sample and taking the maximum value as the peak height. Frequency measurements to perform the FFT were calculated per sample for an entire 96 well plate and plate-wide mean frequencies were used for samples lacking sufficient quality or signal on a given plate. Instability calculations were performed on each distribution of peaks for each identified CAG allele. Peaks were noise-filtered by setting a peak height floor of 50 and unassigning any downstream peaks that were more than 12 base pairs away from the previous peak. Instability was calculated using the PPS metric, transforming the height of every peak of a given allele distribution to a ratio of the main allele peak and summing all expansion-side peaks. After removing duplicates and outliers PPS values for 8,575 Enroll-HD and 654 Registry individuals were generated. Of these, 5,342 had CAG repeats in the range 40-55 units and had whole genome genotyping data and were utilized for blood PPS expansion GWAS.

### ABI PPS somatic expansion GWAS

The blood DNA ABI trace PPS somatic expansion GWAS was carried out with samples having PPS metric, modal expanded *HTT* CAG allele length of 40 to 55 units, who also had age at sample collection of greater than 10 years. This yielded PPS values for 5,342 unique samples. Subsequently, linear regression analysis was performed to calculate the levels of instability adjusted for CAG and age at collection. Briefly, the log transformed PPS data were modeled as a function of CAG repeat length, age at collection, and the age at collection X CAG interaction term. Both CAG length and age at collection were centered around the mean for the regression analysis, generating fitted PPS value for each sample. Sex did not significantly explain variation in the PPS data and therefore was not included in the regression model. Then, the fitted value (i.e., predicted PPS value from the regression model) was subtracted from the actual PPS value for each sample, yielding residual PPS data, which represent the levels of repeat expansion corrected for inherited modal CAG repeat size and age at collection. The residual PPS value was used as the phenotype for the GWAS using a mixed effects model with the same set of covariates that were used for GWAS of clinical phenotypes, (sex, batch of GWAS, first four principal component values, limited to SNVs with MAF > 1%), as described in more detail above in the Methods section.

### Blood somatic expansion ratio phenotype

The somatic expansion ratio (SER = 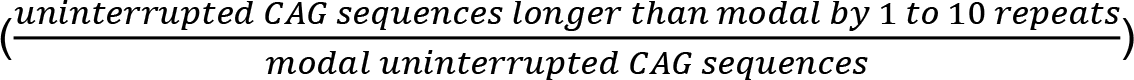 of the *HTT* CAG repeat was quantified by the Glasgow group from the MiSeq read count distributions obtained by alignment of the Q^2^ Solutions MiSeq forward/R1 reads for the 3,999 Enroll-HD participants for which ≥ 250 MiSeq reads could be obtained for the modal expanded CAG length, for which the short *HTT* allele was at least six CAGs shorter than the long *HTT* allele, for which the 40 ≤ modal CAG length ≤ 50 (because inherited CAG length is uncertain above 50 CAGs) and for which age at collection information was available (*i.e.,* exclusion of 580 participants’ samples for which < 250 MiSeq reads were obtained for the modal expanded allele, seven participants’ samples for which the longer CAG allele was ≤ 5 CAGs longer than the shorter CAG allele and both alleles had the same CAG repeat-adjacent sequence, 40 participants’ samples without age at sampling information and 357 participants’ samples with modal long CAG length < 40 and > 50).

Multiple linear regression using the base R function lm was used to derive a CAG and age-independent SER phenotype from the final set of 3,999 MiSeq CAG frequency distributions. These analyses were performed using CAG and age variables centered on their mean. To use the best possible CAG and age adjustment of the SER measure, the following multiple linear regressions models of increasing hierarchical complexity were compared:

SERmodel1 (AIC = -352.5 adjusted r^2^ = 0.841): ln(SER) ∼ *β*_0_ + *β*_1_.CAG + *β*_2_.age + *ε*

SERmodel2 (AIC = -362.2 adjusted r^2^ = 0.843): ln(SER) ∼ *β*_0_ + *β*_1_.CAG + *β*_2_.age + *β*_3_.(CAG x age) + *ε*

SERmodel3 (AIC = -368.6 adjusted r^2^ = 0.845): ln(SER) ∼ *β*_0_ + *β*_1_.CAG + *β*_2_.age + *β*_3_.(CAG x age) + *β*_4_.CAG^2^ + *β*_5_.age^2^ + *ε*

SERmodel4 (AIC = -375.3 adjusted r^2^ = 0.847): ln(SER) ∼ *β*_0_ + *β*_1_.CAG + *β*_2_.age + *β*_3_.(CAG x age) + *β*_4_.CAG^2^ + *β*_5_.age^2^ + *β*_6_.(CAG^2^ x age) + *β*_7_.(CAG x age^2^) + *ε*

SERmodel5 (AIC = -373.3 adjusted r^2^ = 0.847): ln(SER) ∼ *β*_0_ + *β*_1_.CAG + *β*_2_.age + *β*_3_.(CAG x age) + *β*_4_.CAG^2^ + *β*_5_.age^2^ + *β*_6_.(CAG^2^ x age) + *β*_7_.(CAG x age^2^) + *β*_8_.(CAG^2^ x age^2^) + *ε*

Based on model AIC, model 4 was selected for CAG and age adjustments of SER. Although SERmodel4 was associated with a similar adjusted r^2^ to the previously utilized SERmodel1 (Ciosi et al., 2019), it demonstrated a non-linear age effect (*p*_*age*_2 < 0.001) and an interaction CAG-age^2^ (*p*_*CAG* × *age*_^2^ < 0.003) not previously considered. The explanatory variables CCG length, CAG repeat-adjacent sequence and PCR batch (added because they had a small but significant effect on SER variation) were added to SERmodel4 and the residual variation (obtained using the base R residuals function) of this final SERmodel4 (*i.e.,* SERmodel4_final_: ln(SER) ∼ *β*_0_ + *β*_1_.CAG + *β*_2_.age + *β*_3_.(CAG x age) + *β*_4_.CAG^2^ + *β*_5_.age^2^ + *β*_6_.(CAG^2^ x age) + *β*_7_.(CAG x age^2^) + *β*_8_.CCG + *β*_9_.(CAG repeat-adjacent sequence) + *β*_10_.PCRbatch + *ε*) was used as the blood SER phenotype for the GWAS.

### MiSeq blood somatic expansion GWAS

GEMMA (0.98.1) was used to calculate a relatedness matrix for the 3,999 individuals in the somatic expansion GWAS. Association analysis was run using GEMMA (0.98.1) using a linear mixed model with the relatedness matrix, with sex, four principal components from the principal component analysis (PCA) for population structure as covariates.

### HD brain somatic CAG expansion phenotype

The CAG expansion metric for the *HTT* CAG repeat in HD frontal cortex was determined from the MGH method of MiSeq paired-read sequencing using the distribution of uninterrupted CAG repeat sizes. The SER was calculated from the MiSeq read profiles as the number of reads of uninterrupted CAG length longer than modal over relative to the number of reads of modal CAG length. For comparison with blood somatic expansion ratio, the distribution of BA9 cortex expansion ratio values for 493 individuals was calculated as described as model 4 in the blood somatic expansion ratio phenotype section, accounting for CAG size, age at death, CAGXage at death, length of adjacent CCG repeats.

### CAG expansion in cell models

We used our instability R package (https://github.com/zachariahmclean/instability) to process the fragment analysis samples and calculate repeat instability metrics, using a 5% peak threshold and considering peaks 40 repeat units to either side of the modal peak. Since the CAG repeat in these cultures advances as a relatively normally distributed population, we used the average repeat gain, which is the difference between the mean CAG repeat (weighted on the height of the fragment peaks) at four-weeks to day-zero. To statistically compare the groups, we created a linear model that considered the CAG repeat variant and the starting CAG repeat size of the clone, which was used to adjust the average repeat gain based on the starting mean CAG repeat in the plotted data. To do this, we used the coefficient for the effect size of starting repeat length and multiplied that by the difference of each clone to the median starting repeat length, which was then subtracted off the average repeat gain for each clone.

### General procedures for conditional analysis

For a selected candidate region with significant association signals in the final data set, a series of conditional analyses was performed to estimate the number of distinguishable modifier effects in either the age at TFC6 or blood somatic expansion GWAS. To minimize outlier effects, these analyses were carried out for SNVs with MAF >1% except for the inclusion of selected infrequent SNVs at *MSH3*, *FAN1* and *LIG1* that were robustly associated with clinical modification in previous studies or missense variants that emerged in the current study (e.g., rs3218772 at *POLD1*). Briefly, the same statistical model as for single SNV analysis with an additional covariate of the minor allele count of the SNV with the largest absolute effect size in the region of interest was constructed in a mixed effect linear model. If signals p < 1.0E-06 remained after the first conditional analysis, the SNV with the largest absolute effect size (or best p-value when effect sizes did not differ significantly) from that conditional analysis was added as a covariate together with the first SNV and the analysis run again. This sequential conditional analysis was continued until no SNV signals better than p = 1.0E-06 remained. Numbering of modifier effects was arbitrary and did not indicate the order of addition to the sequential conditional analyses.

### Distinguishable clinical modifier effects

For most of the loci identified previously, we confirmed the reported modifier effects in the expanded age at TFC6 data set. We again identified single modifier effects at *MLH1*, *RRM2B* and *CCDC82*. In the *MLH1* region conditioning on rs148648107 (3AM1) left all SNVs with p > 2.0E-03. At *RRM2B*, conditioning on rs79136984 (8AM1) left all other SNVs at p > 8.2E-03. The *CCDC82* region also harbored a single modifier effect because conditioning on rs12365516 (11AM1) left all other SNVs at p > 3.1E-02.

We again found two distinguishable modifier effects in the *PMS1* region. Initially, we conditioned on rs76405401 (2AM2) based on its effect size (0.29), which left rs6434365 (2AM1) as the most significant SNV (conditioned p = 4.90E-13). Conditioning on both rs76405401 and rs6434365 left all SNVs with p > 1.0E-03.

At *MSH3*, we confirmed three modifier effects. We initially conditioned on rs113361582 (5AM2) based on its effect size (0.90), which led to the choice of rs245100 (5AM1; conditioned p = 8.8E-55) for addition to the second round of conditional analysis. The latter identified rs6151716 as the top remaining SNV (conditioned p = 5.8E-17). Conditioning simultaneously on all three SNVs left all other SNVs at p > 1.6E-05.

The *TCERG1* locus revealed two modifier effects. Conditioning on the previously identified rs79727797 (5BM1), which had the largest effect size and most significant p-value, left rs202157262 (5BM2) as the SNV with the top remaining effect size (0.26) and p-value (p = 1.2E-08). Conditioning on both SNVs left all other SNVs at p > 3.3E-05.

In the *PMS2* region, we defined three distinguishable modifier effects. We initially conditioned on rs1805324 (7AM4) based on its effect size (−0.26), which revealed rs1805326 as the genome-wide significant SNV (conditioned p = 1.1E-15) with the largest remaining effect size (0.14). Conditioning simultaneously on both of these tags left rs2345058 as the SNV with the top remained effect size (0.08) and significance (conditioned p = 1.7E-07). Conditioning simultaneously on all three tags left all other SNVs at p > 2.0E-03. Note that the putative 7AM2 modifier effect defined based on the tag rs3779109 in our previous dataset did not survive these sequential conditional analyses, indicating that it was not distinguishable from the combination of the 7AM1, 7AM3 and 7AM4 effects.

At *FAN1*, we conditioned on the strong effect missense variants rs150393409 (15AM1) and rs151322829 (15AM3), which left rs118089305 (15AM5) as the genome-wide significant SNV (conditioned p = 1.1E-12) with the largest absolute effect size (−0.32). Conditioning on all three tags revealed rs3512 (15AM2) as the top remaining SNV (conditioned p = 8.6E-52). Adding rs3512 to the next conditional analysis left all other SNVs at p > 1.1E-04. Note that the modifier effect designated 15AM4 was shown previously to not be distinguishable from the combined effects of the four modifier effects, as confirmed here (Kim et al., 2020).

For *LIG1*, we first conditioned on the strong effect missense variant rs145821638, which pointed to rs3078205 (conditioned p = 5.5E-16). Conditioning on both SNVs left all other SNVs at p > 8.8E-05. The previously implicated 19AM2 effect did not survive these analyses indicating that it is not distinguishable from the combined effects of 19AM1 and 19AM3.

At *POLD1*, we conditioned on the top SNV rs112837068 (19BM1) to discover the missense variant rs3218772 as tagging a second effect (19BM2, conditioned p = 7.5E-07). Conditioning on both left all other SNVs at p > 2.5E-02.

At *MED15*, we conditioned on the top SNV rs177425 (22AM1) to find all SNVs left at p > 1.8E-04, indicating a single distinguishable modifier effect.

At the new locus detected by the 22BM1 modifier effect in the age at TMSq75 analysis, conditioning on the tag SNV rs8137409 in these data left all signals at the locus at p >1.2E-02, again indicating a single distinguishable modifier effect.

### Conditional analysis of non-canonical repeats

Among study subjects with age at motor onset, DNA samples for 7,553 participants were also subjected to sequencing analysis of the repeat, revealing individuals carrying expanded CAG repeats with canonical (n = 7,285), CAACAG-dup (n = 120), CAA/CCA-loss (n = 92), CAA-loss (n = 29), or CCA-loss (n = 27) alleles. To determine the impacts of non-canonical repeats on motor onset and to identify their tagging-SNVs, we analyzed each variant in the 5 Mb region of chromosome 4p in a mixed effects model. Initially, the regression model was corrected by the same set of covariates that were used in the GWAS plus all non-canonical repeat alleles (i.e., CAACAG-dup, CAA/CCA-loss, CAA-loss, CCA-loss). Conditioning all non-canonical repeat alleles left all SNV signals at p > 4.9E-06. To test a specific non-canonical repeat allele, the variable for that allele was excluded from the covariates, and the results were compared to the data corrected for all non-canonical repeat alleles. For example, to evaluate the impact of CAACAG-dup alleles, dummy variables for CAA/CCA-loss, CAA-loss, and CCA-loss were included in the regression analysis model. This analysis revealed significant SNVs, which represent tagging-SNVs for CAACAG-dup chromosomes. Similarly, analysis without correction for CAA/CCA-loss showed significant SNVs, identifying SNVs tagging CAA/CCA-loss chromosomes. In contrast, the regression models without CAA-loss or CCA-loss left all signals at p > 5.9E-06, suggesting a lack of tagging variants and/or no significant impact on modifying motor onset.

### Somatic expansion modifier effects

In the somatic expansion GWAS, we conditioned on chromosome 2 on rs113983130 (2ABEM1) to define rs79939438 (conditioned p = 4.9E-11) as the tag for 2ABEM2. Conditioning on both left all SNVs (MAF >1%) at p > 3.3E-04.

At *HTT*, conditioning on the top SNV, rs146151652 (tagging 4ABEM1), left all other SNVs with MAF > 1% at p > 1.5E-06.

At *MSH3*, despite having an MAF <1%, we conditioned first on rs113361582 (5ABEM2) because of its robust effect in the clinical data and genome-wide significant p-value and strong effect (−0.22) in the somatic expansion GWAS. This pointed to rs1650689 (5ABEM3, conditioned p = 9.3E-19) as the top remaining SNV. Conditioning on both SNVs then revealed rs245105 (5ABEM1, conditioned p = 1.1E-10). Conditioning simultaneously on all three SNVs left all remaining SNVs with MAF > 1% at p > 2.5E-05.

In the *PMS2* region, we found a set of five distinguishable modifier effects. We initially conditioned on rs1805324 (7ABEM1) based on its high effect size (0.10), which revealed rs12673868 (7ABEM2) and rs75973354 (7ABEM3) as SNVs of comparable MAF (6.4% and 4.4% respectively) and significant p-value (conditioned p = 6.9E-09 and 1.4E-08). Conditioning separately on each of these two SNVs revealed that neither eliminated but instead increased the signal from the other, making them clearly distinguishable. Consequently, we next conditioned simultaneously on rs1805324, rs12673868, and rs75973354, which raised the significance of an infrequent missense SNV, rs63750123 (7ABEM4), beyond our threshold (conditioned p = 1.4 E-07). Adding this SNV to the other three for conditional analysis then revealed rs2079819 (7ABEM5) as the top remaining SNV (conditioned p = 3.5E-07). Simultaneously conditioning on all five tags left the remaining SNVs with MAF >1% at p > 5.4E-04.

At *MLH3,* the top signals and effect sizes were both associated with very frequent SNVs. Conditioning on the top SNV, rs5809690, left all other > 1% MAF SNVs at p > 4.8E-03, suggesting a single modifier effect (14ABEM1).

At *FAN1,* conditioning on the strong effect missense variants rs150393409 (15ABEM1) and rs151322829 (15ABEM3), revealed rs118089305 (15ABEM4, conditioned p = 5.7E-07) as the remaining SNV with the highest effect size (0.07). Conditioning on all three tags identified rs61260387 (15ABEM2) as the top remaining SNV (conditioned p = 7.3E-37). Conditioning on all four SNVs left the rest (MAF > 1%) at p > 9.3E-06.

The chromosome 17 locus encompassing *ATAD5, ADAP2,* and *TEFM*, showed rs117662433 (17ABEM1) as a genome-wide significant SNV (p = 4.9E-09), with lower MAF (4.3%) and higher effect size (0.058) than the top SNV (rs62070643 tagging 17ABEM2, p = 1.0E-14, MAF 26.4%, effect 0.035). Conditioning on rs117662433 revealed rs8067252 as the top remaining SNV (conditioned p = 6.0E-10). Conditioning on both of these SNVs left all other MAF >1% SNVs at p >7.6E-04.

### LD calculation

SNVs that tag the haplotype marked by the top SNV in the association and/or conditional analysis were identified based on LD r^2^ values, which were estimated from the clinical sample data by using the PLINK program. In the association region plots, we set the r^2^ square threshold as 0.8 to mark SNVs in high LD with the lead SNV.

### LD MED15 CAG and chr 22AM1 SNV

*MED15* CAG repeat number was extracted from SPARK whole genome sequence CRAM files for 491 unrelated European ancestry individuals. Reads aligned to exon 7 of *MED15* (hg38: chr22:20,566,467-20,566,817) were selected and the CAG repeat number was obtained from each read by selecting the nucleotide sequence between a start motif of CAACAGCAGCAG and a stop motif of GCTTTGCAG. Reads sharing the same sequence were aggregated and the top two most frequent sequences were selected. The sample was classified as a heterozygous genotype if the second most frequent sequence has a 25% or greater proportion of the top two aggregates. If two frequent sequences were not observed, the most frequent sequence was classified as a homozygous genotype. The length of the uninterrupted CAG repeat of these aggregated sequences was measured, thereby determining CAG repeat number. Variant calls from the SPARK whole genome sequencing VCF file were also utilized to determine the 22AM1 top tagging SNV rs177425 [hg38 chr22:20573256:C:T] alleles. The alternate allele had an 17% MAF in this set of 491 unrelated individuals, similar to the MAF of 19% in TOPMED and gnomAD databases. The two main *MED15* CAG repeat alleles were CAG12 (reference) and CAG13. The following haplotypes were observed: five CAG9 all phased with the SNV reference allele; three CAG10 of which one CAG10 phased with SNV reference allele and two with the alternate allele; eight CAG11 of which six phased with SNV reference allele and two phased with the alternate allele; 795 CAG12 of which 751 phased with SNV reference allele and 44 phased with the alternate allele; 163 CAG13 of which 50 phased with SNV reference allele and 113 phased with the alternate allele; six CAG14 of which four phased with SNV reference allele and two with the alternate allele; one CAG15 which phased with the alternate SNV allele and one CAG16 which phased with the SNV reference allele. For the purposes of calculating D’, the relatively infrequent CAG repeat haplotypes were grouped with the CAG12 reference haplotypes (called CAG12+) and compared with the CAG13 haplotypes. This yielded 768 CAG12+ with the SNV reference allele (χ^11^= 0.78); 51 CAG12 with the SNV alternate allele (χ^21^= 0.05); 50 CAG13 with the SNV reference allele (χ^12^ =0.05); and 113 CAG13 with the SNV alternate allele (χ^22^ =0.12). The allele frequencies were: SNV reference 0.83=p1; SNV alternate=0.17=p2; CAG12 reference =0.83=q1; CAG13 alternate = 0.17=q2. D’ and square of correlation between the alleles at the two loci (r^2^) were then calculated using: D = (χ^11^)(χ^22^)-(χ^12^)(χ^21^) yields D=0.09 and, since Dmax is smaller of p1q2 or p2q1, therefore D’=D/Dmax= 0.63 and r^2^=D^2^/p1p2q1q2 = 0.40.

### Pathway analysis

The same pathways were used as in (Bellenguez et al., 2022). The assignment of Gene Ontology (GO) terms to human genes was obtained from the “gene2go” file, downloaded from NCBI (ftp://ftp.ncbi.nlm.nih.gov/gene/DATA/) on March 11^th^ 2020. “Parent” GO terms were assigned to genes using the ontology file downloaded from the Gene Ontology website (http://geneontology.org/docs/download-ontology/) on the same date. GO terms were assigned to genes based on experimental or curated evidence of a specific type, so evidence codes IEA (electronic annotation), NAS (non-traceable author statement), RCA (inferred from reviewed computational analysis) excluded. Pathways were downloaded from the Reactome website (https://reactome.org/download-data) on April 26^th^ 2020. Biocarta, KEGG and Pathway Interaction Database (PID) pathways were downloaded from v7.1 (March 2020) of the Molecular Signatures Database (Subramanian et al., 2005) (https://www.gsea-msigdb.org/gsea/msigdb/index.jsp). Analysis was restricted to GO terms containing between 10 and 2000 genes. No size restrictions were placed on the other gene sets, since there were many fewer of them. This resulted in a total of 10,043 gene sets for analysis.

Gene set enrichment analyses were performed in MAGMA (de Leeuw et al., 2015), correcting for the number of SNVs in each gene, linkage disequilibrium (LD) between SNVs and LD between genes. The measure of pathway enrichment is the MAGMA “competitive” test (where the association statistic for genes in the pathway is compared to those of all other protein-coding genes). We used the “mean” test statistic, which uses the sum of -log(SNP p-value) across all genes as the association statistic for genes. The primary analysis assigned variants to genes if they lie within the gene boundaries, but a secondary analysis used a window of 35kb upstream and 10kb downstream to assign SNPs to genes, as in (Network Pathway Analysis Subgroup of Psychiatric Genomics Consortium, 2015).

### GWAS quality control and data management - MGH

1. Quality controls parameters of imputed SNPs were generated using the following options by the PLINK program.

plink --file GENOTYPE_DATA --freq plink --bfile GENOTYPE_DATA --missing

plink --bfile GENOTYPE_DATA --hardy --nonfounders --hwe 0

Then, imputed SNPs were subjected to quality controls as described in the method section.

2. Construction of kinship matrix was based on the following code using genotype file in the PLINK format in the GEMMA program.

gemma -bfile GENOTYPE_DATA -gk 1

3. Association analysis using a linear mixed effect model was based on following code.

GENOTYPE_DATA file set in the PLINK file format included the test phenotypes. gemma -bfile GENOTYPE_DATA -k KINSHIP_MATRIX -c COVARIATE -lmm 1 -maf 0

4. Conditional analyses of selected regions were based on the following options, which include a set of covariates and the genotype of conditioning SNPs.

gemma -bfile GENOTYPE_DATA -k KINSHIP_MATRIX -c COVARIATE+GENOTYPE -lmm 1 -

maf 0

5. LD of a test variant with other SNVs were calculated using the PLINK program.

plink1.9 --bfile GENOTYPE_DATA --r2 --ld-snp TEST_VARIANT --ld-window-kb 5000 --ld-window 999999 --ld-window-r2 0

6. Regional plots were generated by R program using plot (X, Y, options) command.

### GWAS quality control and data management - Glasgow

1. Pre-imputation QC

plink --bfile GENOTYPE_FILE_TO_BE_IMPUTED --mind 0.05 --maf 0.01 --geno 0.05 --make- bed

Data formatting to match reference panel (TOPMed)

Calculate allele frequencies

plink1.9 --bfile GENOTYPE_FILE_QC --freq

perl ../program/HRC-1000G-check-bim.pl -b GENOTYPE_FILE_QC -f FREQUENCY_FILE -r

../program/PASS.Variantsbravo-dbsnp-all.tab.gz -h

./Run-plink.sh

for ((i=1; i< 23; i++)) do

bcftools sort GENOTYPE_FILE$i.vcf -Oz done

1. Post-imputation processing Merge vcf chromosome files

bcftools concat chr{1..22}.dose.vcf.gz -Oz

Filter vcf file for imputation quality score (RSq) > 0.5

bcftools view -i ’R2>.5’ -Oz IMPUTED_VCF_FILE > IMPUTED_VCF_FILE Convert vcf to binary plink format

plink --vcf IMPUTED_VCF_FILE --make-bed --keep-allele-order --double-id Filter for minor allele frequency

plink --bfile GENOTYPE_FILE --maf 0.01 --make-bed

1. Construction of kinship matrix for somatic expansion GWAS: gemma -bfile GENOTYPE_DATA -gk 1

1. Association analysis of somatic expansion ratio:

gemma -bfile GENOTYPE_DATA -k KINSHIP_MATRIX -c COVARIATES_FILE -lmm 1 -maf 0

## Supplemental Information

**Figure S1.**
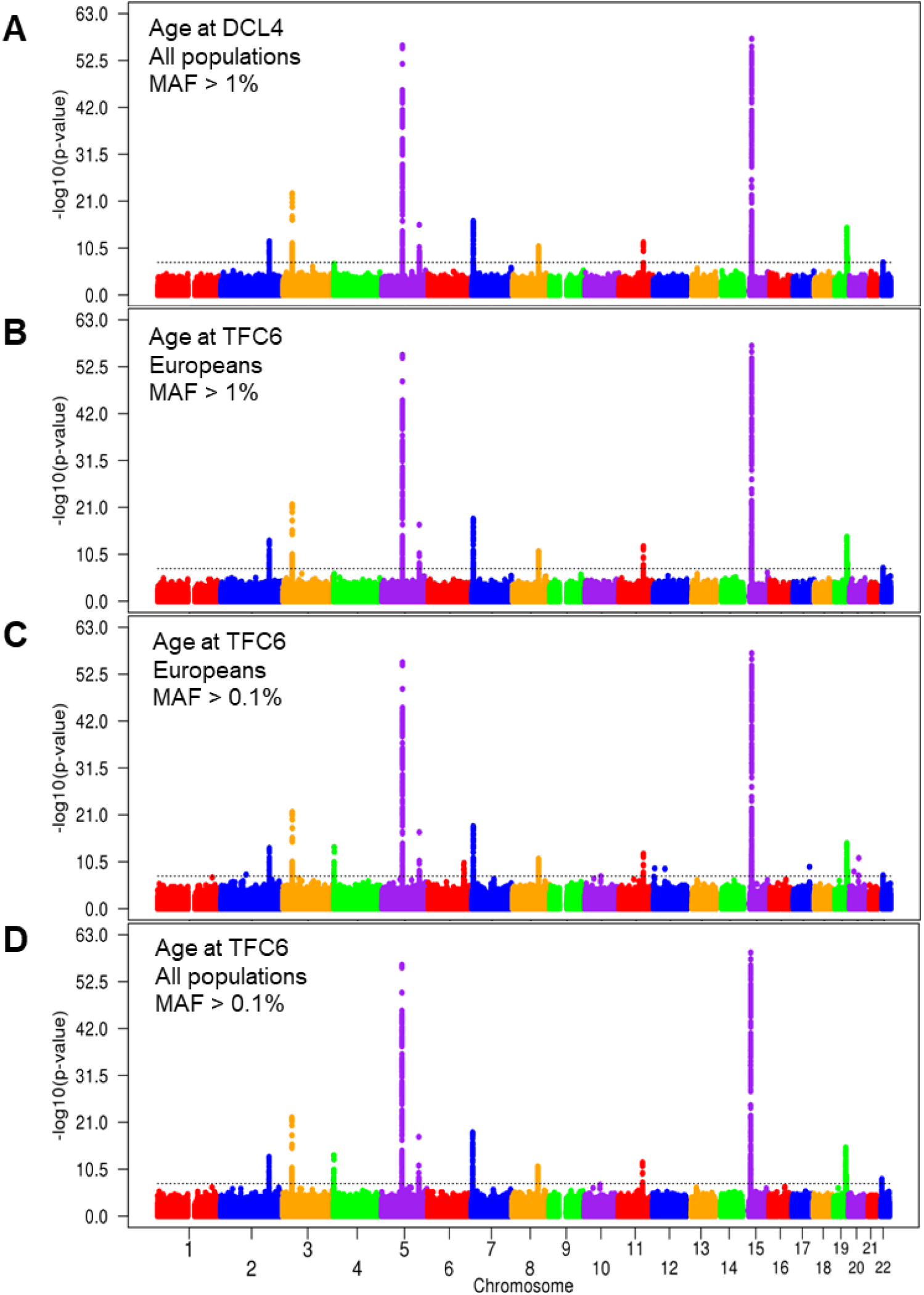
– Comparison of GWAS results across age at DCL4, ancestry and low MAF. For comparison with each other and with Figure 1A, GWAS results are plotted with each point representing a test SNV and significance shown as -log10(p-value). The dotted black line represents the threshold for genome-wide significance (p = 5.0E-08). **A**. GWAS of algorithmically predicted age at DCL4 in all populations (n = 11,408) at SNV MAF >1%. **B**. GWAS of age at TFC6 only in individuals of European ancestry (n = 11,185) at SNV MAF > 1%. **C**. GWAS of age at TFC6 only in individuals of European ancestry at SNV MAF > 0.1%. **D**. GWAS of age at TFC6 in all individuals regardless of ancestry (n = 11,698) at SNV MAF > 0.1%. The overall pattern of results did not differ substantially across these analyses except for infrequent SNVs (MAF < 1%) in Europeans. Although a few infrequent SNVs have previously detected strong, reproducible, significant effects at established loci with multiple modifier haplotypes such as *MSH3*, *FAN1*, and *LIG1* and at *HTT*, most isolated genome-wide significant modifier signals have proved inconsistent across previous studies. We proposed that infrequent SNVs can produce spurious signals when present in a few phenotypic outliers. In the current GWAS, with few exceptions, infrequent SNVs that newly achieved significance in the European-only analysis (panel C) had a higher reported MAF in African-ancestry individuals. These SNVs were no longer significant after the inclusion of the non-Europeans suggesting that they were due to phenotypic outliers among the Europeans. Beyond the infrequent SNVs reported previously at *MSH3*, *FAN1* and *LIG1* and at *HTT* (Table S1, Figure S4), the only infrequent SNVs that were genome-wide significant in the all-population GWAS (rs145090839, rs193001369, rs17864672, rs824382) occurred with age at motor onset. Yet, none of these achieved even suggestive significance (p < 1.0E-05) with either age at DCL4 or age at TFC6, supporting the notion that they are likely due to motor onset outliers.

**Figure S2.**
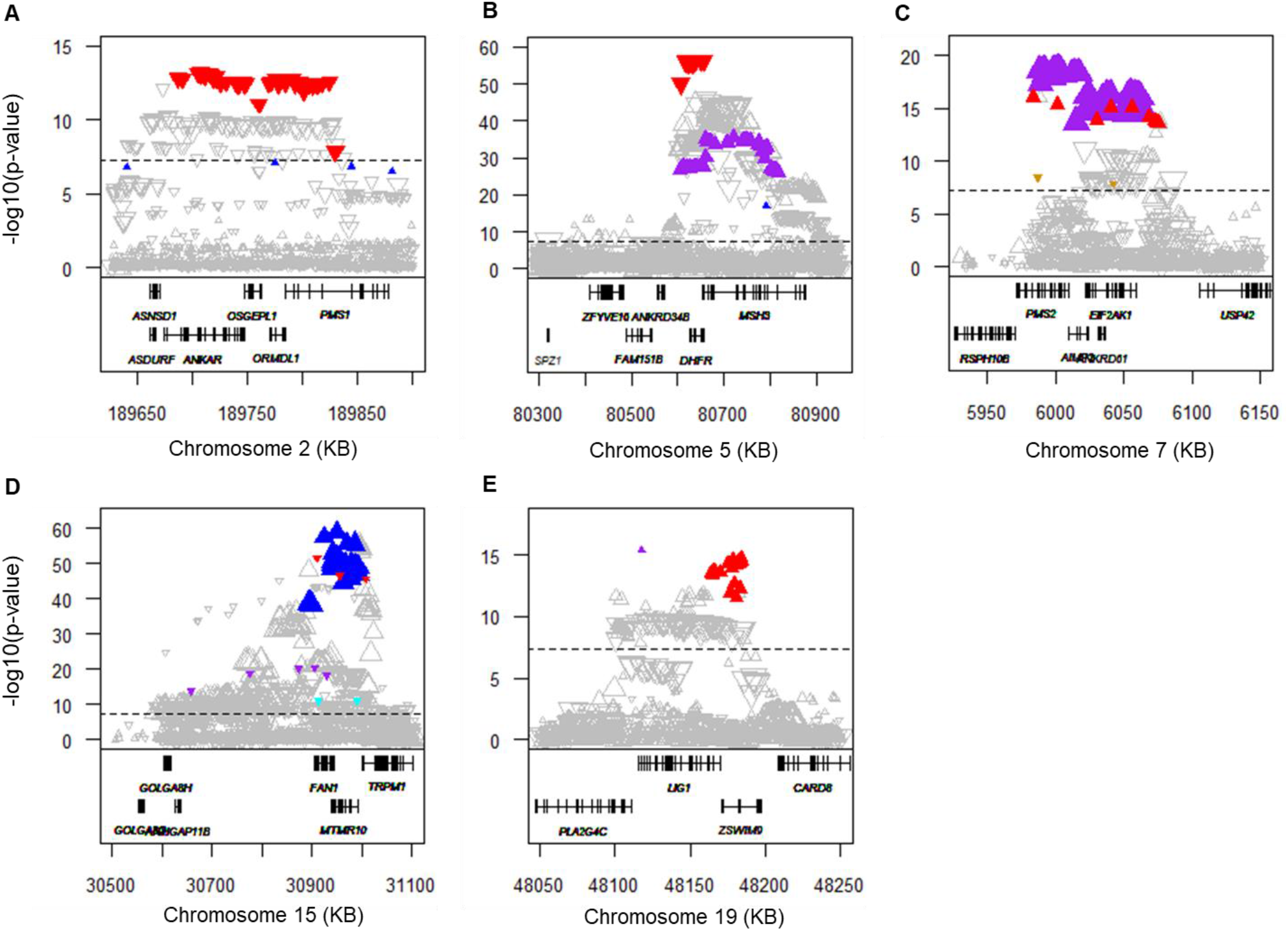
– Loci with multiple clinical modifier effects. Association results for age at TFC6 from Figure 1A are shown in the regions of *PMS1* (**A**), *MSH3* (**B**), *PMS2* (**C**), *FAN1* (**D**), and *LIG1* (**E**), with genes in each region shown below the plot. The dashed black line represents the threshold for genome-wide significance (p = 5.0E-08). For each distinguishable modifier effect (see Table S1), the top SNV and all other SNVs showing r^2^ > 0.8 with the top SNV are shown as filled triangles (red for 2AM1, 5AM1, 7AM1, 15AM1 and 19AM1; blue for 2AM2, 5AM2, and 15AM2; purple for 5AM3, 7AM3, 15AM3 and 19AM3, brown for 7AM4; and, cyan for 7AM5 and 15AM5) while all other SNVs are represented by unfilled triangles. The triangle’s size and orientation reflect its SNV MAF and direction of clinical effect (upward pointing = delaying; downward pointing = hastening).

**Figure S3.**
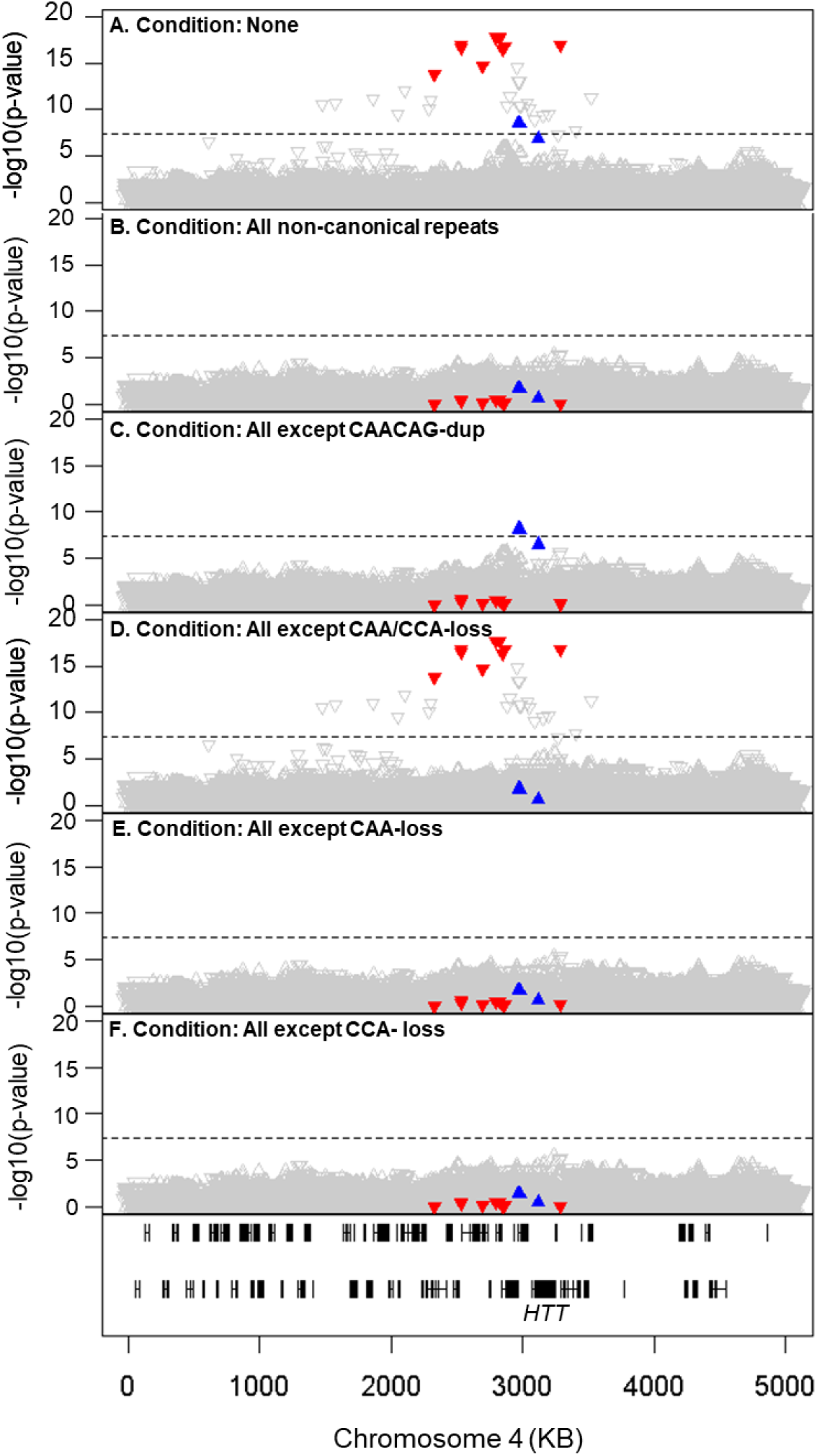
– Infrequent SNVs capture modifier effects of non-canonical *HTT* CAG repeat alleles. **A**. For all HD participants with *HTT* alleles defined by sequence analysis (n =7,553), age at motor onset GWAS results are plotted for all SNVs > 0.1% MAF in the terminal 5 Mb of chromosome 4p16.3, containing *HTT*. The dashed line in this and subsequent panels represents the threshold for genome-wide significance (p = 5.0E-08). Each SNV is represented by a triangle whose size and orientation reflect its MAF and direction of effect (upward pointing = delaying; downward pointing = hastening). For each of two significant modifier effects, the top SNV and all other SNVs showing r^2^ >0.8 with the top SNV are shown as filled triangles: red = motor onset-hastening effect; blue = motor onset-delaying effect. [Because the age at TFC6 (and all other algorithmically predicted landmarks) prediction dataset was based on CAG genotyping in the clinical studies and not on sequencing, the algorithmically predicted phenotypes could not be analyzed without bias in this region due to differences with sequence-determined CAG lengths for non-canonical alleles.] **B**. Association results for age at motor onset are plotted as in panel A, after conditioning on all non-canonical sequences, which removed all significant signals. **C**. Association results after conditioning on all non-canonical sequences except the CAACAG-dup allele, revealing surviving signal for SNVs shown as blue triangles that tag HD chromosomes with the CAACAG-dup sequence. **D**. Association results after conditioning on all non-canonical sequences except the CAA/CCA-loss allele, revealing surviving signal for SNVs shown as red triangles that tag HD chromosomes with the CAA/CCA-loss sequence. Conditioning on all non-canonical alleles except CAA-loss (**E**) or CCA-loss (**F**) removed all significant SNV signals.

**Figure S4.**
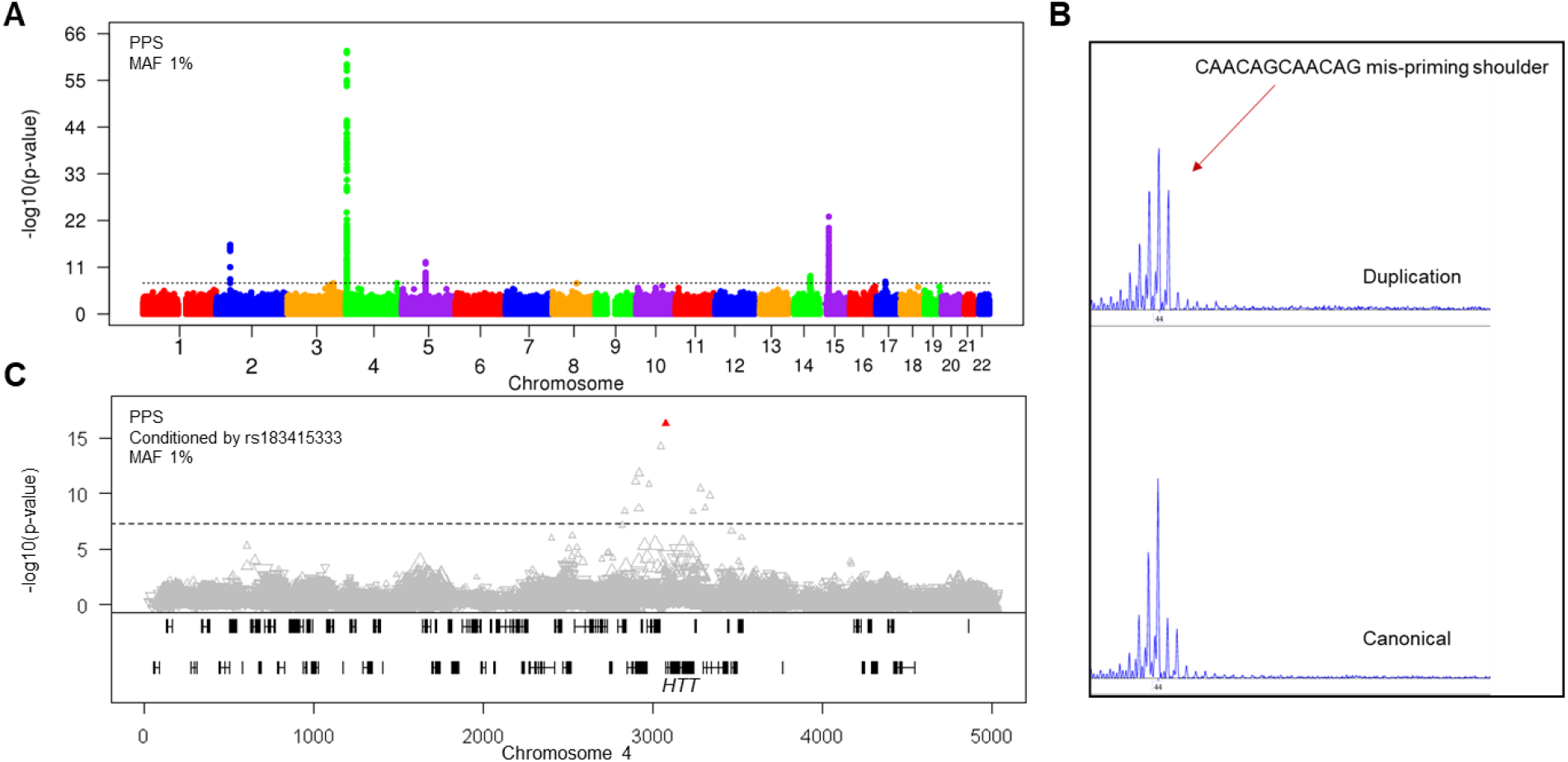
– GWAS of somatic CAG expansion based on the Peak Proportional Sum (PPS) Method from ABI fragment sizing traces. **A**. GWAS of somatic CAG expansion measured by the PPS method applied to ABI blood CAG sizing traces is shown for SNVs with MAF > 1%. The dashed black line represents the threshold for genome-wide significance (p = 5.0E-08). **B**. An example of ABI traces from a CAACAG-dup allele (top) and a canonical allele (bottom), both carrying uninterrupted repeat lengths of 44 CAGs. The top panel reveals a mispriming artefact that scores as CAG expansion in the PPS method, leading to spurious signal at *HTT* in the GWAS of panel A. **C**. Association analysis of the PPS CAG expansion phenotype conditioned on rs183415333, which tags the non-canonical CAACAG-dup allele on HD chromosomes, is shown for SNVs with MAF > 1% in the *HTT* region. Conditioning removes the spurious signal and leaves rs146151652 as the top SNV, the same *HTT* 5’-UTR SNV detected in the MiSeq somatic CAG expansion GWAS (Figure 3B, Table S2).

**Figure S5.**
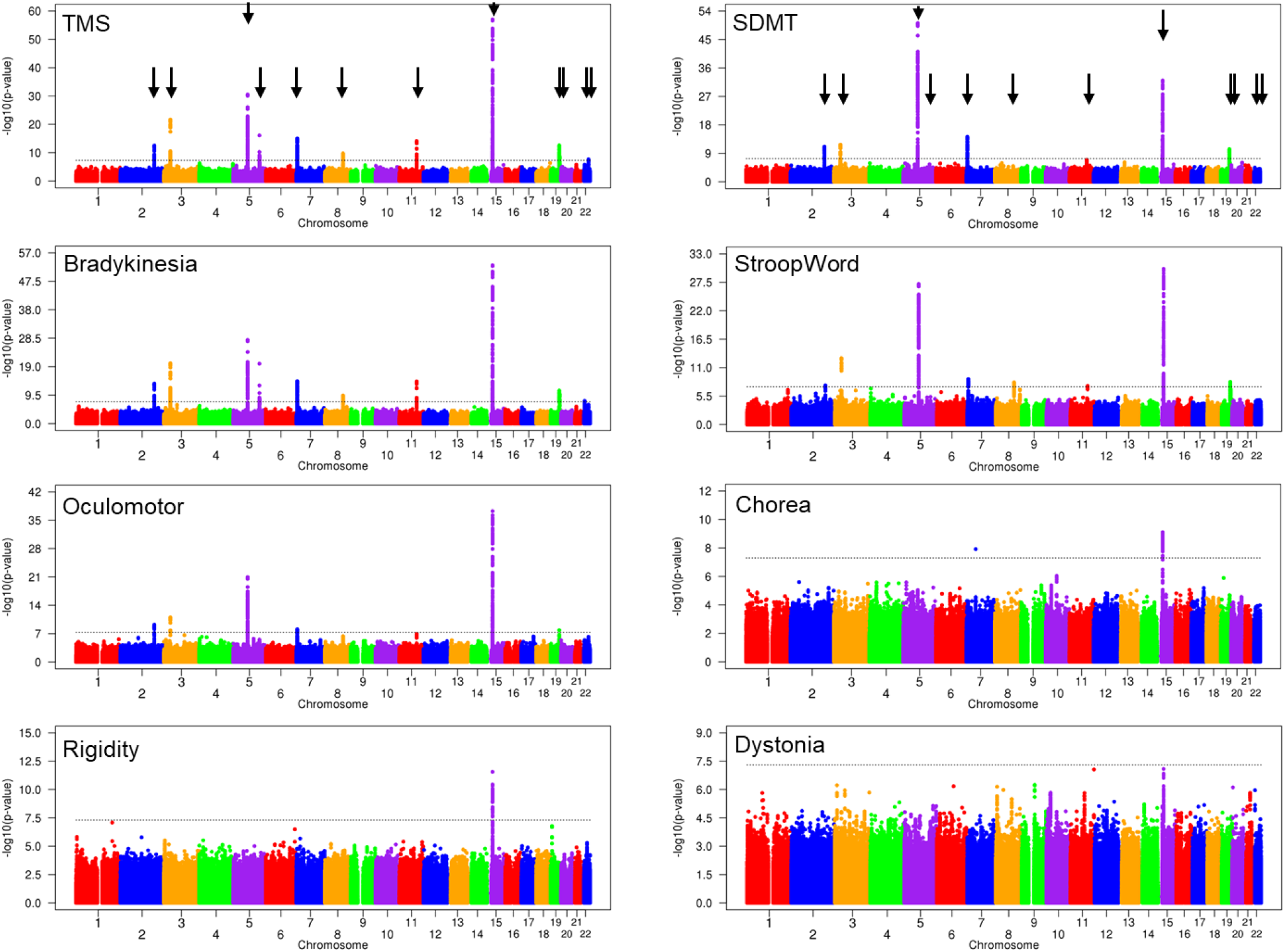
– Comparison of GWAS for late-disease clinical landmarks. Separate GWAS at MAF > 1% are shown for each of the late-disease algorithmically predicted landmarks based on the 25% quantile for the Symbol Digit Modalities Test (SDMT) and the StroopWord Reading Test (StroopWord) and the 75% quantile for UHDRS-Total Motor Score (TMS) and each of its subcomponents, bradykinesia, oculomotor, chorea, rigidity and dystonia. For ease of comparison, vertical arrows are placed in the top panels at the positions of any locus showing genome-wide significant signal in any clinical phenotype analysis. The dashed black lines represent the threshold for genome-wide significance (p = 5.0E-08).

**Table S1.** – Modifiers of HD clinical landmarks

| Chromosome | Modifier effect<sup>1</sup> | Candidate gene | Tag variant | Minor allele | MAF | SNV # | Beta | P value | Triangle color in Figures |
| --- | --- | --- | --- | --- | --- | --- | --- | --- | --- |
| 2 | 2AM1 | *PMS1* | chr2:189707695:T:C | C | 20.9 | rs6434365 | -0.12 | 5.85E-14 | red |
|  | 2AM2 |  | chr2:189775136:C:T | T | 1.3 | rs76405401 | 0.29 | 7.73E-08 | blue |
| 3 | 3AM1 | *MLH1* | chr3:37076854:G:A | A | 30.5 | rs148648107 | 0.14 | 8.01E-23 |  |
| 5 | 5AM1 | *MSH3* | chr5:80637274:A:G | G | 24.9 | rs245100 | -0.23 | 5.63E-57 | red |
|  | 5AM2 |  | chr5:80790685:A:G | G | 0.3 | rs113361582 | 0.90 | 9.14E-18 | blue |
|  | 5AM3 |  | chr5:80720482:GCTT:G | G | 16.0 | rs6151716 | 0.22 | 3.36E-36 | purple |
| 5 | 5BM1 | *TCERG1* | chr5:146507273:G:A | A | 2.7 | rs79727797 | 0.32 | 1.83E-18 | red |
|  | 5BM2 |  | chr5:146473061:AGGAAG:A | A | 2.0 | rs202157262 | 0.25 | 3.29E-08 | blue |
| 7 | 7AM1 | *PMS2* | chr7:5982995:C:T | T | 14.8 | rs1805326 | 0.15 | 1.06E-16 | red |
|  | 7AM3 |  | chr7:6001000:A:C | C | 40.9 | rs2345058 | 0.12 | 1.82E-19 | purple |
|  | 7AM4 |  | chr7:5986899:C:T | T | 2.1 | rs1805324 | -0.26 | 3.47E-09 | brown |
| 8 | 8AM1 | *RRM2B* | chr8:102201412:G:T | T | 7.8 | rs79136984 | -0.16 | 7.88E-12 |  |
| 11 | 11AM1 | *CCDC82* | chr11:96389264:C:T | T | 19.6 | rs12365516 | 0.11 | 9.99E-13 |  |
| 14 | 14AM1<sup>2</sup> | *MLH3* | chr14:75140342:G:A | A | 45.8 | rs175438 | -0.06 | 3.71E-06 | red |
| 15 | 15AM1 | *FAN1* | chr15:30910758:G:A | A | 1.2 | rs150393409 | -0.87 | 2.69E-52 | red |
|  | 15AM2 |  | chr15:30942802:G:C | C | 31.1 | rs3512 | 0.22 | 1.08E-53 | blue |
|  | 15AM3 |  | chr15:30905792:C:T | T | 0.8 | rs151322829 | -0.64 | 3.02E-21 | purple |
|  | 15AM5 |  | chr15:30912434:C:T | T | 1.8 | rs118089305 | -0.31 | 8.63E-12 | cyan |
| 19 | 19AM1 | *LIG1* | chr19:48183794:ACC:A | A | 13.0 | rs3078205 | 0.15 | 1.59E-15 | red |
|  | 19AM3 |  | chr19:48117686:C:A | A | 0.2 | rs145821638 | 1.24 | 4.46E-16 | purple |
| 19 | 19BM1 | *POLD1* | chr19:50397291:T:C | C | 9.7 | rs112837068 | 0.13 | 6.62E-10 | red |
|  | 19BM2 |  | chr19:50398939:C:T | T | 1.0 | rs3218772 | 0.28 | 3.43E-06 | blue |
| 22 | 22AM1<sup>3</sup> | *MED15* | chr22:20573256:C:T | T | 21.3 | rs177425 | -0.09 | 5.03E-09 | red |
|  |  |  | chr22:20573515:G:A | A | 21.3 | rs165666 | -0.09 | 5.03E-09 |  |
| 22 | 22BM1 | *CCDC134,* | chr22:41963789:G:A | A | 11.0 | rs8137409 | 0.12 | 2.25E-08 | red |
|  |  | *SREBF2,* |  |  |  |  |  |  |  |
|  |  | *SHISA8,* |  |  |  |  |  |  |  |
|  |  | *TNFRSF13C,* |  |  |  |  |  |  |  |
|  |  | *CENPM,* |  |  |  |  |  |  |  |
|  |  | *SMIM45,* |  |  |  |  |  |  |  |
|  |  | *SEPTIN3* |  |  |  |  |  |  |  |

<sup>1</sup> All effects from age at TFC6 GWAS except 22BM1 from age at TMSq75 GWAS
<sup>2</sup> This putative effect did not achieve genome-wide significance but is listed for comparison with somatic expansion GWAS
<sup>3</sup> Two 'top' SNVs had identical significance and effect size

**Table S2.**
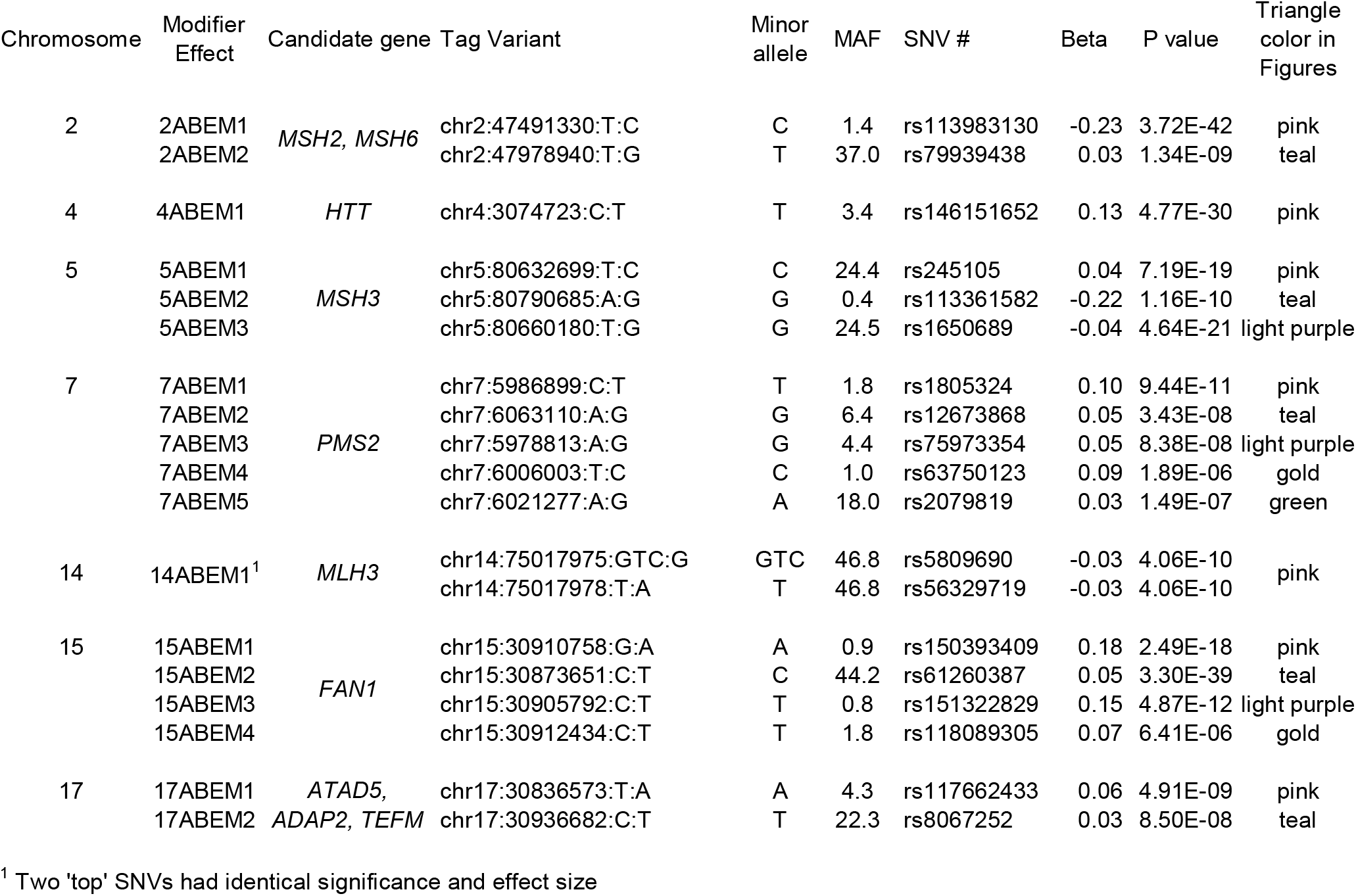
– Modifiers of somatic CAG expansion in blood DNA

